# Environmental heterogeneity and microbial community architecture shape N_2_O fluxes in intertidal coastal sediments

**DOI:** 10.64898/2026.11.19.767851

**Authors:** Marion Dolivet-Maréchal, Carlos Palacin-Lizarbe, Guillaume Schwob, Henri M.P. Siljanen, Dhiraj Paul, Abigaïl Delort, Jonathan Gervaix, Charline Creuzé des Châtelliers, Sabine Schmidt, Mathis Cognat, David Sebag, Olivier Taugourdeau, Clara Schübert, Nathalie Labourdette, Isabelle Bertrand, Lorenzo Rossi, Xavier Le Roux, Agnès Richaume, Alessandro Florio

## Abstract

Seagrass meadows are key “blue carbon” ecosystems, yet their contribution to nitrous oxide (N_2_O) emissions remains poorly constrained. We investigated environmental and microbial descriptors of N_2_O fluxes in intertidal sediments of Arcachon Bay, home to Europe’s largest *Zostera noltii* meadow. At six sites, we measured *in situ* N_2_O fluxes in adjacent bare and vegetated sediments and characterized sediment properties, N_2_O-cycling genes using qPCR and targeted metagenomics, and microbial co-occurrence networks. Fluxes ranged from −3.7 to 8.4 µmol N_2_O m^−2^ d^−1^. Spatial location was the strongest descriptor of flux variation, with inner-bay sites showing higher emissions associated with environmental conditions, carbon accumulation, pH, and microbial community characteristics. Vegetation effects were comparatively small and site-dependent. Metagenomic descriptors of N_2_O-cycling genes were more informative than qPCR-based descriptors. Taxa associated with *nirS* (*Thiohalophilus*), *nirK* (*Planktotalea*) and *nosZ1* (*Roseobacter*) were identified as potential contributors to N_2_O production or reduction, respectively. These findings indicate that spatially structured environmental conditions and microbial community characteristics better describe N_2_O flux variation than seagrass presence alone. Measurements across tidal and seasonal cycles are needed to establish the temporal consistency of these patterns.

## 1. INTRODUCTION

Seagrass meadows are globally important “blue carbon” ecosystems that store large amounts of organic carbon (C_org_) in their sediments over long timescales, thereby contributing to climate change mitigation^1,2^. Although they occupy only ∼0.1% of the ocean surface, seagrass meadows account for up to 18% of marine carbon (C) burial^1,3^. However, their net climatic benefit also depends on greenhouse gases (GHGs) exchanges with the atmosphere, which can offset the cooling effect of CO_2_ uptake through C burial^4,5^. Among these gases, nitrous oxide (N_2_O) is a potent GHG with a global warming potential approximately 273 times higher than CO_2_ over a 100-year timescale^6^ and a major contributor to stratospheric ozone depletion^7,8^. Recent estimates suggest that N_2_O emissions from seagrass sediments can offset up to 24.7 ± 0.3% of sedimentary C burial^9^, highlighting the need to better understand the processes controlling N_2_O dynamics in coastal vegetated ecosystems.

N_2_O fluxes from seagrass sediments result from the balance between multiple microbially mediated nitrogen (N) transformations, including nitrification, denitrification, nitrifier denitrification and dissimilatory nitrate reduction to ammonium (DNRA). The transformations are catalysed by phylogenetically diverse microbial communities carrying key functional genes^10–12^. Seagrass sediments exhibit steep redox gradients over millimetre scales, with oxygenated surface layers and rhizosphere microzones surrounding largely anoxic bulk sediments^13,14^. These contrasting microhabitats promote tight coupling between aerobic and anaerobic N transformations, making seagrass sediments potential hotspots for both N_2_O production and consumption^5,15^. Key functional genes involved in these processes include *nirK* and *nirS*, encoding nitrite reductases involved in heterotrophic denitrification, which represents the dominant pathway of N_2_O production in coastal sediments^11,16^, and *nosZ* clades I and II, encoding nitrous oxide reductases responsible for the final denitrification step through the reduction of N_2_O to N_2_^17–20^. In seagrass sediments, *nosZ* clade II communities are often highly diverse and widespread, including members of the Gammaproteobacteria, Alphaproteobacteria and Bacteroidetes as well as non-denitrifying N_2_O reducers such as Anaeromyxobacter dehalogenans, highlighting their potential importance in regulating net N_2_O emissions^21^.

The balance between N_2_O production and consumption is controlled by the interaction between environmental conditions and microbial community structure. Sediment organic matter (OM) content and quality, oxygen availability and dissolved inorganic nitrogen (DIN) concentrations are recognized as major drivers of N_2_O dynamics in coastal sediments^11^. However, beyond the abundance of individual functional genes, increasing evidences suggest that microbial diversity, community composition and ecological interactions may influence N_2_O fluxes on an ecosystem-scale^22,23^. Microbial co-occurrence networks provide a complementary framework to functional gene approaches, by revealing potential ecological associations and patterns of organization^24,25^. Although recent studies have examined microbial networks in seagrass sediments and marine N_2_O cycling, the role of microbial network architecture in regulating N_2_O production and reduction in coastal vegetated sediments remains poorly understood^26,27^.

Empirical observations of N_2_O fluxes from seagrass ecosystems are scarce and highly variable. Global syntheses indicate that seagrass meadows may act as either weak sources or sinks of N_2_O depending on species identity, environmental conditions, and methodological approaches^10,28^. Most studies rely on laboratory incubations or submerged water–sediment flux measurements, whereas *in situ* sediment–air N_2_O exchange in intertidal seagrass ecosystems remains largely unexplored^4,10,28^. Moreover, the effect of seagrass vegetation itself is still debated. While some studies report enhanced N_2_O emissions in vegetated sediments^15^, others suggest that seagrass may stimulate N_2_O reduction through rhizosphere processes, including the supply of labile C to denitrifying microorganisms^29–31^, or by limiting N availability for N_2_O production^21,32^. These contrasting observations highlight the need to identify the main abiotic and microbial descriptors explaining N_2_O variability in seagrass ecosystems.

Temperate dwarf seagrass species such as *Zostera noltii*, which dominates intertidal habitats in north-western Europe, are particularly underrepresented in N_2_O studies despite their extensive distribution and sediment C _org_ storage capacity^28^. Previous work in estuarine systems, including that of Dai et al.^33^, showed that denitrifiers (*nirS*) dominate high-N environments, whereas ammonia oxidizers (*amoA*) prevail under low-N conditions, while *nosZ* clade II is widespread and highly diverse. These findings suggest that the composition of the microbial community, together with environmental conditions, plays a key role in regulating N_2_O emissions. However, no studies have simultaneously investigated vegetated and adjacent bare sediments while integrating N_2_O flux measurements with the composition, functional potential and co-occurence networks of microbial communities.

In Arcachon Bay (Figure 1), which hosts the largest *Z. noltii* population in Europe, we previously demonstrated that N_2_O fluxes were higher in vegetated than in bare sediments at two of the six studied sites, with total emissions offsetting 24.7 ± 0.3% of local C burial^9^. However, the spatial variability of these emissions and their microbial drivers across the bay remain unresolved. Building on this work, the present study aims to (i) investigate the spatial patterns of N_2_O fluxes across Arcachon Bay, including both vegetated and bare sediments, (ii) determine how site location and seagrass presence influence the composition, functional potential and network architecture of N_2_O-cycling microbial communities, and (iii) identify the abiotic and biotic descriptors explaining the variability of N_2_O fluxes in intertidal seagrass sediments. We hypothesised that N_2_O fluxes would be primarily driven by sediment N stocks, and that enhanced emissions in seagrass meadows would be associated with an increased abundance and diversity of N_2_O-producing communities (*nirK* and *nirS*).

**Figure 1.**
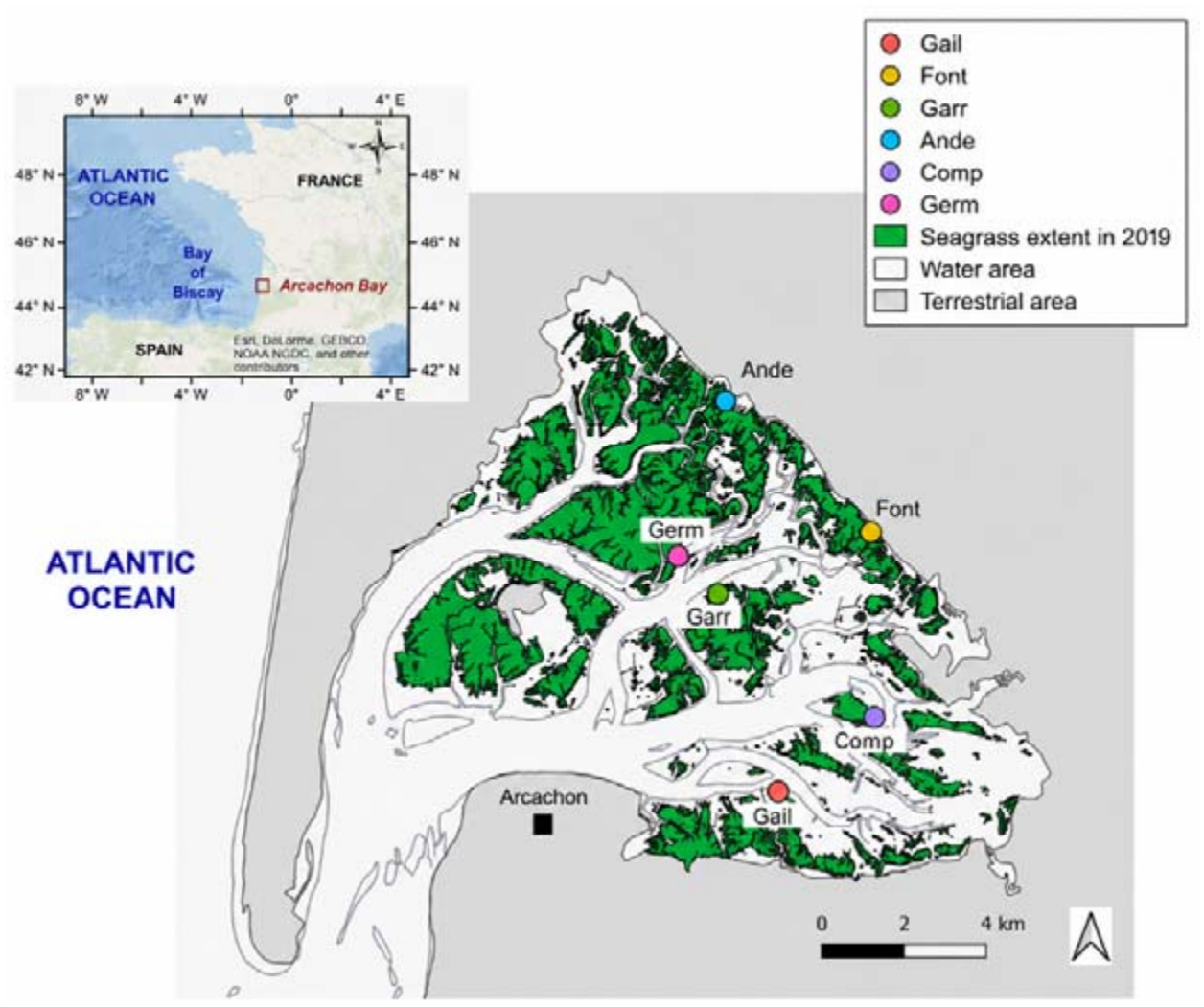
Location of Arcachon Bay and the six studied sites with 2019 *Z. noltii* coverage.

To test these hypotheses, we combined *in situ* N_2_O flux measurements conducted across Arcachon Bay during daytime at low tide, allowing standardized comparisons of peak emissions among sites^9^, with sediment physicochemical characterization, qPCR and targeted metagenomics of N_2_O-cycling genes (*nirK, nirS, nosZ1* and *nosZ2*). Microbial co-occurrence networks were reconstructed to assess potential ecological associations and community organization, while mixed linear models (MLMs) and random forest analyses (RFA) were used to identify the main abiotic and biotic descriptors of N_2_O variability. Understanding the controls of N_2_O emissions in *Z. noltii* meadows provides new insights into the microbial regulation of N-cycling and helps to refine assessments of the climate benefits of coastal blue C ecosystems.

## 2. MATERIALS AND METHODS

### 2.1. Study site and experimental design

Arcachon Bay is a temperate coastal lagoon located on the French southern Atlantic coast (44°40⍰ N, 1°10⍰ W), covering approximately 174 km^2^ (Figure 1). The bay hosts the largest European population of the dwarf seagrass *Z. noltii*, which forms extensive intertidal meadows on sheltered mudflats^34^. Over the past three decades, their coverage has declined by ∼44%, largely due to extreme weather events such as heatwaves and storms^34–36^.

To encompass the environmental variability of Arcachon Bay, six sites were selected based on contrasting hydrodynamic conditions, sediment characteristics and vegetation status (Figure 1). Each site included adjacent vegetated (*Z. noltii*) and unvegetated (“Seagrass” and “Bare”, respectively) sediments, with six replicates per subsite (see Dolivet-Maréchal et al.^9^ for further details). Historical aerial imagery analyses indicate that these bare areas were vegetated prior to 2005. Both types of habitat have remained environmentally stable for over 20 years, providing a consistent framework for comparison. Two field campaigns were conducted during the weeks commencing September 12, 2022, and August 28, 2023, i.e. during the peak biomass of *Z. noltii*^*9,34*^. In 2022, sediment cores were collected to characterise C burial and screen for potential GHG production. In 2023, GHG fluxes (N_2_O, CH_4_ and CO_2_) were measured *in situ* using opaque static flux chambers, deployed at midday and at low tide, following a standardised layout (5-meter radius). Immediately after each flux measurement, three sediment cores (10 cm depth) were collected at the chamber location and pooled into a single composite sample per replicate, representing the surface sediment layer commonly used for GHG and microbial analyses. In total, of 71 sediment samples with 62 N_2_O flux measurements (four to six flux measurements per subsite) were analysed across the bay. To preserve *in situ* conditions and prevent microbial alteration, the samples were stored in the dark at either 4°C (for incubations and sediment characterizations) or −20°C (for molecular analyses). Sediment physico-chemical parameters were quantified as described in the Supplementary materials.

### 2.2. *In situ* N_2_O measurements

N_2_O fluxes were measured using a LI-COR static Smart Chamber system (model 8200-01S) coupled with a trace gas analyzer (LI-7820; LI-COR Biosciences, USA). The chamber was deployed on pre-installed PVC collars at each replicate location, and gas concentrations were monitored over a 3-min period after stabilization. Data was recorded every second and processed using LI-COR SoilFluxPro− software. Fluxes were calculated from temporal changes in N_2_O concentration and expressed as µmol N_2_O m^−2^ d^−1^ following Dolivet-Maréchal et al.,^9^.

### 2.3. Abundance of bacterial 16S rRNA and N_2_O-producing and -consuming functional genes

The abundances of bacterial 16S rRNA and N_2_O-producing (*nirS, nirK*) and -consuming (*nosZ1, nosZ2*) functional genes, were estimated *via* quantitative real-time PCR after DNA extraction and quantification from sediment samples (see full methodological details in Supplementary Materials and Methods and Table S1). To better capture the combined potential of N_2_O-producing and N_2_O-consuming microbial communities, the abundances of *nirK* and *nirS* were summed to create the variable *nirKS*, while the abundances of *nosZ1* and *nosZ2* were summed to create the variable *nosZ12*. In addition, a proxy of the balance between N_2_O production and consumption potential was calculated as a qPCR-based ratio (ratio_qPCR), defined as the sum of *nir* gene abundances (*nirS* + *nirK*) divided by the sum of *nosZ* gene abundances (*nosZ1* + *nosZ2*).

### 2.4. Targeted metagenomic sequencing and functional gene diversity

A targeted metagenomic approach was used to characterize the diversity and composition of N-cycling microbial communities. Sequence capture by probe hybridization was performed to enrich 12 functional genes involved in mineral N transformations (*nirS, nirK, nosZ1, nosZ2, amoA, nrfA, norB, hdh, nxrB, nifH, napA* and *narG*) following Putkinen et al. and Siljanen et al.^37,38^. Sequencing data were processed using a dedicated bioinformatic pipeline described in Supplementary Methods. Relative abundances of functional genes and taxa were converted into absolute abundances using qPCR-derived gene copy numbers. For *nirS, nirK, nosZ1*, and *nosZ2* genes, the 30 most abundant taxa across all samples were retained for downstream analyses. Gene-specific richness and Shannon diversity were calculated from phylogenetic placements and are referred to as gene richness (*gene*_R) and gene diversity (*gene*_D), respectively. Ratios between N_2_O-producing and N_2_O-consuming gene diversity metrics were also calculated (Ratio_R and Ratio_D). All sequencing data are available in the NCBI Sequence Read Archive under accession number PRJNA1261814.

### 2.5. Co-occurrence network

Microbial co-occurrence networks were constructed from taxa carrying *nirS, nirK, nosZ1*, and *nosZ2* genes to compare network architectures between sampling locations (Inner vs. Outer bay) and vegetation types (Bare vs. Seagrass). Network inference, topological analyses, node role classification, and robustness analyses followed the workflow described by Schwob et al.^39^. Full details of the network construction and analysis are provided in the Supplementary Methods.

### 2.6. Random forest analysis and mixed linear models

To assess the environmental and microbial drivers of N_2_O fluxes, RFA and MLMs were employed following the approach described in Dolivet-Maréchal et al.^40^. Briefly, explanatory variables were grouped into predefined subsets according to their nature (Table S2A–I), and separate models were constructed to identify the most relevant descriptors for each subset. Random Forest models were first applied separately to each subset as a complementary non-parametric approach to rank variable importance, identify potential nonlinear relationships, and pre-select the most relevant descriptors before MLM analyses (full details in Supplementary Methods). Due to computational constraints, only the 30 most abundant taxa per functional gene (*nirK, nirS, nosZ1* and *nosZ2*) were included in the RFA analyses. The ten most important taxonomic descriptors identified from the RFA analyses of the four functional genes (*nirK, nirS, nosZ1* and *nosZ2*) were retained for subsequent MLM analyses.

For MLM analyses, explanatory variables were transformed when required and scaled using z-scores to allow comparison of their relative effects. Model selection was performed using the dredge() function from the *MuMIn* package, retaining only models including significant descriptors (p < 0.05). To avoid overfitting and improve model interpretability, the number of explanatory variables was limited to a maximum of five per model. Model performance was assessed using the corrected Akaike information criterion (AICc) and R^2^ values. Linear mixed-effects models were fitted using the lme() function in R (v4.3.1), testing random intercepts for spatial variability among sampling sites, habitat types (Seagrass vs. Bare sediments), their combination (subsites) and the location in the bay (Inner vs. Outer). Nested random-effect structures were also evaluated, including habitat type nested within site and habitat type nested within bay location. Fixed-effect models using generalized least squares (gls()) were also tested to allow comparison of model performance.

The most relevant descriptors identified from each subset-specific analysis were merged into a combined dataset. RFA and MLM were then reapplied to this combined dataset to identify the key environmental and microbial descriptors of N_2_O fluxes and evaluate their overall explanatory power.

### 2.7. Statistical analyses

All data analyses were conducted using R (version 4.5.2). Results are reported as means ± standard error (SE). Variables were log_10_- or square root-transformed when necessary to meet analysis assumptions and reduce the influence of extreme values. Differences between bare and vegetated sediments were tested by site and across the entire bay, and differences between Inner and Outer sites, were assessed using Wilcoxon tests. Spearman’s rank correlation tests were used to examine relationships between N_2_O fluxes and all studied variables. Only significant correlations are reported, with significance levels indicated as follows: # for p < 0.1, * for p < 0.05, ** for p < 0.01, and *** for p < 0.001. Correlation coefficients (ρ) are represented using a color scale, with red indicating negative correlations, bleu indicating positive correlations, and the intensity of the color reflecting the strength of the correlation. Principal Component Analysis (PCA) was performed to explore patterns among physical, chemical, microbial, and GHG variables across all sampling sites, using the *FactoMineR* package in R. Among the network metrics, only heterogeneity_location was included in the PCA because it was the most informative network property in the N_2_O flux models.

## 3. RESULTS

### 3.1. Spatial environmental heterogeneity and N_**2**_**O fluxes across Arcachon Bay**

PCA revealed a marked spatial organization of the sampling sites across Arcachon Bay (Figures 2 and S1a,b). The first principal component (PC1, 30% of the total variance) separated the two Inner sites (Andernos and Fontaines), located closer to the terrestrial zone, from the four Outer sites, influenced by oceanic conditions. PC2 explained an additional 16% of the variance. Inner sites were located at an average distance of 19.4 ± 0.1 km from the open ocean, whereas the Outer sites were situated at an average distance of 15.7 ± 0.5 km (Figure 2b). Based on this clear separation, subsequent analyses distinguished Inner (Ande and Font) and Outer (Germ, Garr, Comp and Gail) sites.

**Figure 2.**
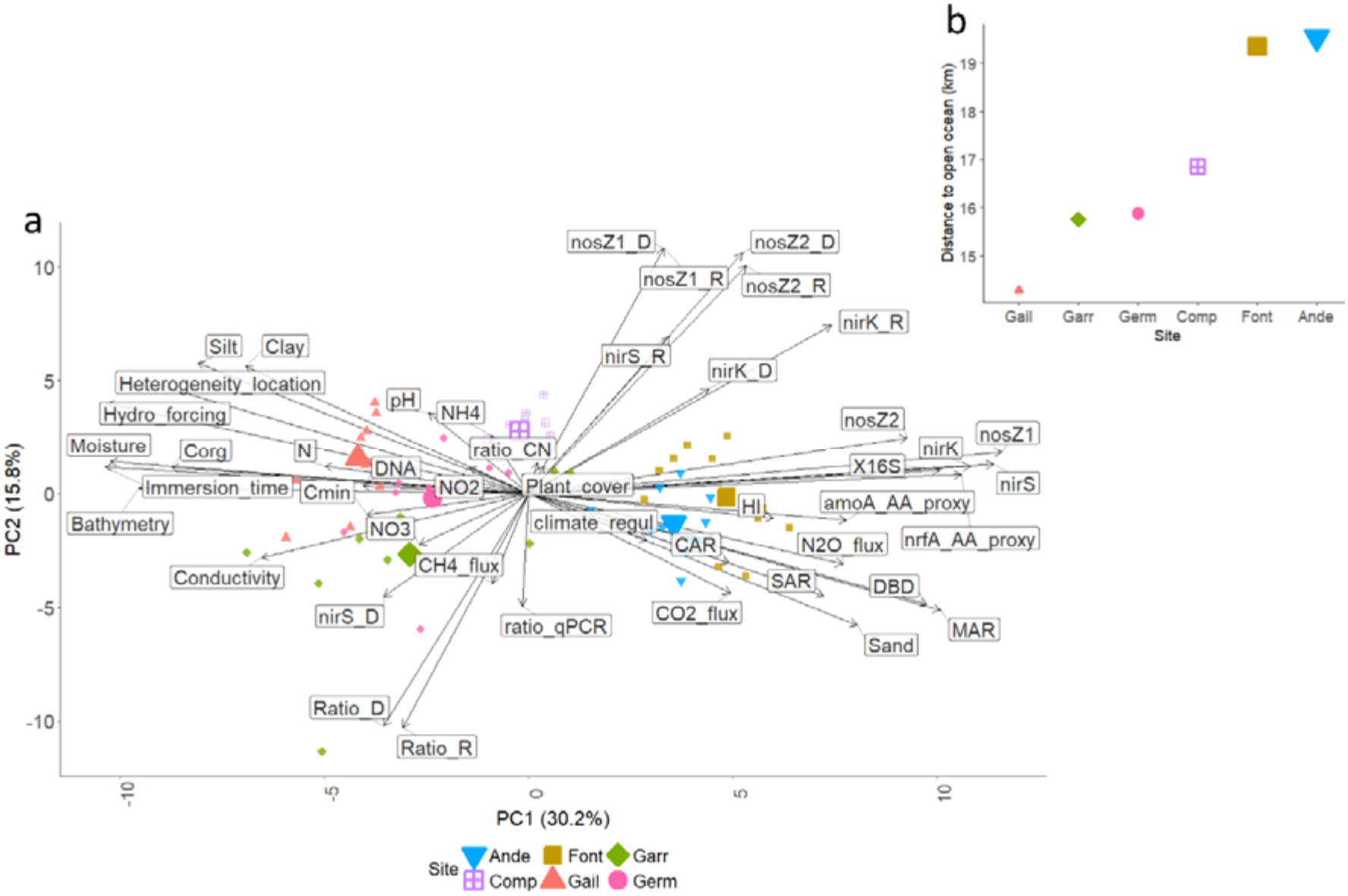
Principal component analysis (PCA) biplot illustrating the relationships among physical, chemical, microbial, GHG, and microbial co-occurrence network (heterogeneity_location) variables across all sampling sites. (a) PCA biplot showing variable loadings on the first two principal components and individual samples (n = 62, after excluding samples with missing data). Arrows represent variable loadings; vector lengths were rescaled by a constant factor for visualization while preserving their relative magnitudes. Point shapes and colours indicate site identity, and larger symbols represent site barycentres. (b) Distance of each site from the open ocean (km); symbol size represents the relative position of each site along the PC1 gradient. Variable abbreviations are defined in the Methods.

Inner sites (on the positive side of the PC1) were associated with higher labile OM, as reflected by the hydrogen index (HI), sedimentation accumulation rate (SAR), carbon accumulation rate (CAR), and coarser sandy sediments (Tables 1 & S3). These sites also exhibited higher abundances of total prokaryotes (16S rRNA gene copies) and N-cycling genes, including nitrification (*amoA*), N_2_O production (*nirS* and *nirK*), N_2_O reduction (*nosZ1* and *nosZ2*), DNRA (*nrfA*) and greater richness and diversity of *nirK* and *nosZ2*. Overall, the summed abundance of N-cycling genes was positively correlated with HI (Spearman’s ρ = 0.56, p < 3.08 × 10^−7^) and negatively correlated with total sediment C_org_ content (ρ = −0.49, p = 1.41 × 10^−5^) (Figure S2). In contrast, Outer sites (on the negative side of the PC1) were characterized by finer, deeper and wetter sediments, stronger hydrodynamic forcing, longer immersion times, and higher total N and DIN (NH_4_^+^ and NO_2_^-^) (Figure 2 and Table S3). Although vegetated and bare sediments showed some overlap in the PCA ordination, vegetation did not produce a clear separation of samples (Figure S1c), indicating that site location was the primary driver of environmental and microbial variability across the bay.

**Table 1.**
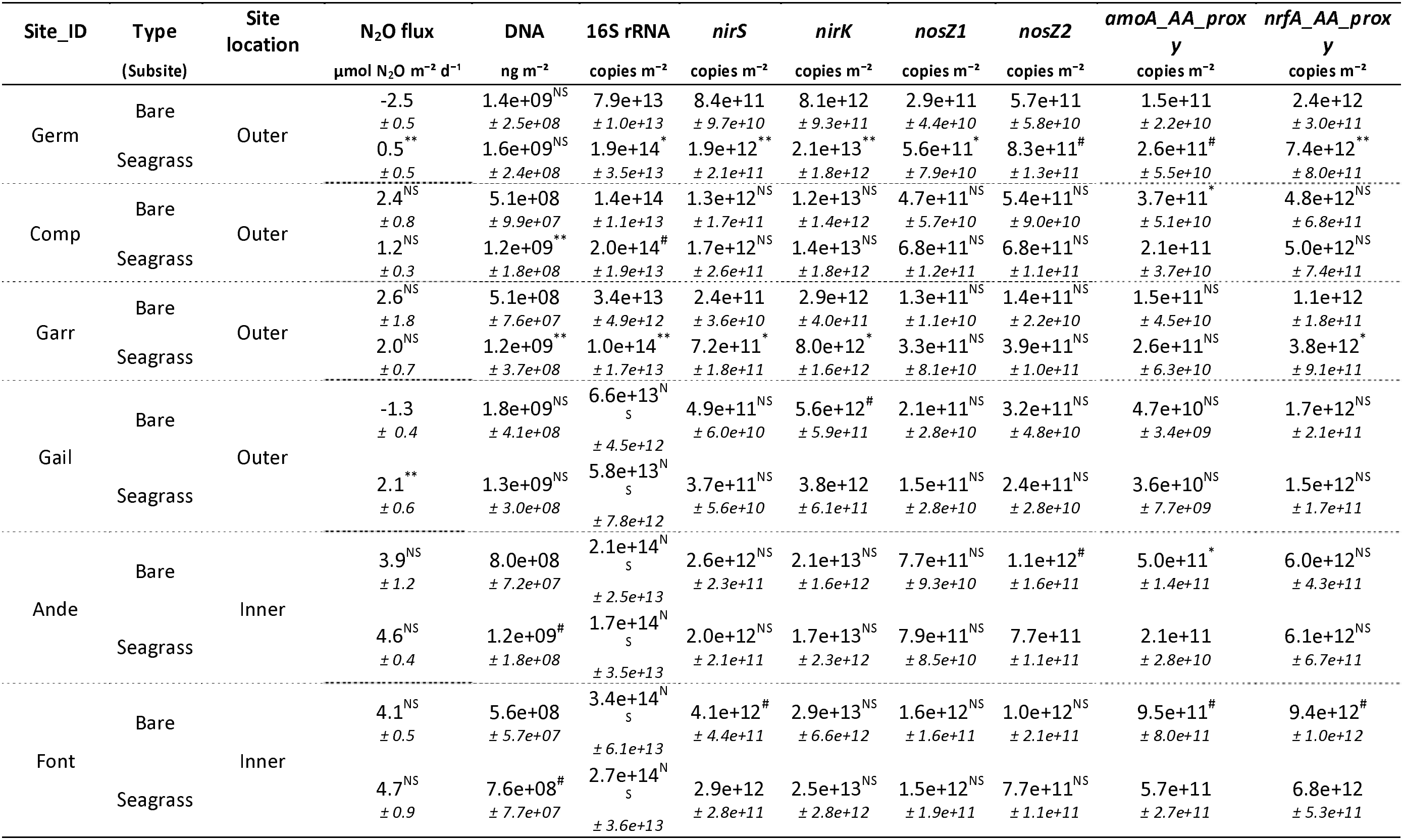
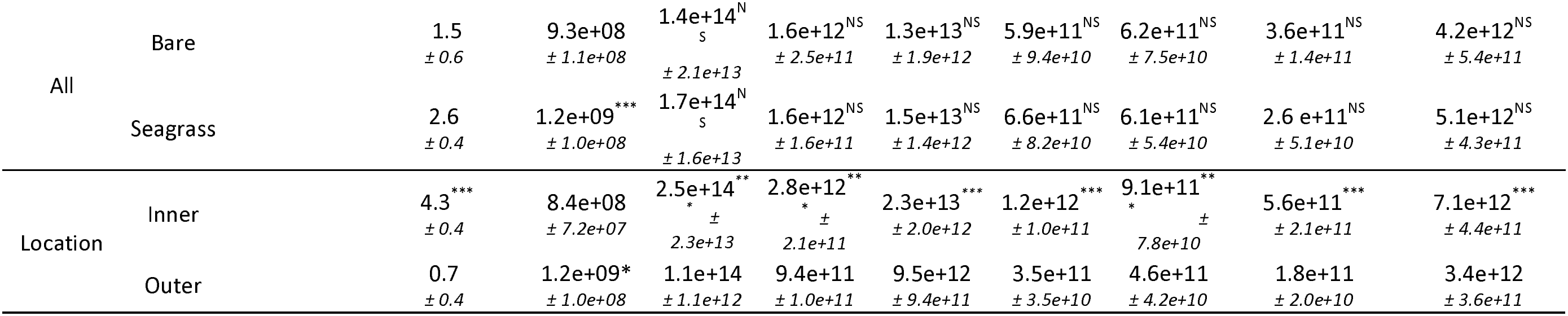
GHG-related variables and microbial data for the studied sites, grouped by sediment type: Bare or vegetated (Seagrass). Sites are ered by increasing N_2_O fluxes measured in seagrass sediments. Significant differences (Wilcoxon tests) between bare and vegetated iments or Inner and Outer sites are indicated as follows: # (p < 0.1), * (p < 0.05), ** (p < 0.01) ), *** (p < 0.001), and NS (not significant). mple sizes were n = 71 for microbial measurements and n = 62 for N_2_O fluxes.

| **Site_ID** | **Type** | **Site location** | **N<sub>2</sub>O flux** | **DNA** | **16S rRNA** | ***nirS*** | ***nirK*** | ***nosZ1*** | ***nosZ2*** | ***amoA_AA_proxy*** | ***nrfA_AA_proxy*** |
|---|---|---|---|---|---|---|---|---|---|---|---|
|  | **(Subsite)** |  | **µmol N<sub>2</sub>O m<sup>-2</sup> d<sup>-1</sup>** | **ng m<sup>-2</sup>** | **copies m<sup>-2</sup>** | **copies m<sup>-2</sup>** | **copies m<sup>-2</sup>** | **copies m<sup>-2</sup>** | **copies m<sup>-2</sup>** | **copies m<sup>-2</sup>** | **copies m<sup>-2</sup>** |
| Germ | Bare | Outer | -2.5<br>*± 0.5* | 1.4e+09<sup>NS</sup><br>*± 2.5e+08* | 7.9e+13<br>*± 1.0e+13* | 8.4e+11<br>*± 9.7e+10* | 8.1e+12<br>*± 9.3e+11* | 2.9e+11<br>*± 4.4e+10* | 5.7e+11<br>*± 5.8e+10* | 1.5e+11<br>*± 2.2e+10* | 2.4e+12<br>*± 3.0e+11* |
|  | Seagrass |  | 0.5<sup>\*\*</sup><br>*± 0.5* | 1.6e+09<sup>NS</sup><br>*± 2.4e+08* | 1.9e+14<sup>\*</sup><br>*± 3.5e+13* | 1.9e+12<sup>\*\*</sup><br>*± 2.1e+11* | 2.1e+13<sup>\*\*</sup><br>*± 1.8e+12* | 5.6e+11<sup>\*</sup><br>*± 7.9e+10* | 8.3e+11<sup>#</sup><br>*± 1.3e+11* | 2.6e+11<sup>#</sup><br>*± 5.5e+10* | 7.4e+12<sup>\*\*</sup><br>*± 8.0e+11* |
| Comp | Bare | Outer | 2.4<sup>NS</sup><br>*± 0.8* | 5.1e+08<br>*± 9.9e+07* | 1.4e+14<br>*± 1.1e+13* | 1.3e+12<sup>NS</sup><br>*± 1.7e+11* | 1.2e+13<sup>NS</sup><br>*± 1.4e+12* | 4.7e+11<sup>NS</sup><br>*± 5.7e+10* | 5.4e+11<sup>NS</sup><br>*± 9.0e+10* | 3.7e+11<sup>\*</sup><br>*± 5.1e+10* | 4.8e+12<sup>NS</sup><br>*± 6.8e+11* |
|  | Seagrass |  | 1.2<sup>NS</sup><br>*± 0.3* | 1.2e+09<sup>\*\*</sup><br>*± 1.8e+08* | 2.0e+14<sup>#</sup><br>*± 1.9e+13* | 1.7e+12<sup>NS</sup><br>*± 2.6e+11* | 1.4e+13<sup>NS</sup><br>*± 1.8e+12* | 6.8e+11<sup>NS</sup><br>*± 1.2e+11* | 6.8e+11<sup>NS</sup><br>*± 1.1e+11* | 2.1e+11<br>*± 3.7e+10* | 5.0e+12<sup>NS</sup><br>*± 7.4e+11* |
| Garr | Bare | Outer | 2.6<sup>NS</sup><br>*± 1.8* | 5.1e+08<br>*± 7.6e+07* | 3.4e+13<br>*± 4.9e+12* | 2.4e+11<br>*± 3.6e+10* | 2.9e+12<br>*± 4.0e+11* | 1.3e+11<sup>NS</sup><br>*± 1.1e+10* | 1.4e+11<sup>NS</sup><br>*± 2.2e+10* | 1.5e+11<sup>NS</sup><br>*± 4.5e+10* | 1.1e+12<br>*± 1.8e+11* |
|  | Seagrass |  | 2.0<sup>NS</sup><br>*± 0.7* | 1.2e+09<sup>\*\*</sup><br>*± 3.7e+08* | 1.0e+14<sup>\*\*</sup><br>*± 1.7e+13* | 7.2e+11<sup>\*</sup><br>*± 1.8e+11* | 8.0e+12<sup>\*</sup><br>*± 1.6e+12* | 3.3e+11<sup>NS</sup><br>*± 8.1e+10* | 3.9e+11<sup>NS</sup><br>*± 1.0e+11* | 2.6e+11<sup>NS</sup><br>*± 6.3e+10* | 3.8e+12<sup>\*</sup><br>*± 9.1e+11* |
| Gail | Bare | Outer | -1.3<br>*± 0.4* | 1.8e+09<sup>NS</sup><br>*± 4.1e+08* | 6.6e+13<sup>NS</sup><br>*± 4.5e+12* | 4.9e+11<sup>NS</sup><br>*± 6.0e+10* | 5.6e+12<sup>#</sup><br>*± 5.9e+11* | 2.1e+11<sup>NS</sup><br>*± 2.8e+10* | 3.2e+11<sup>NS</sup><br>*± 4.8e+10* | 4.7e+10<sup>NS</sup><br>*± 3.4e+09* | 1.7e+12<sup>NS</sup><br>*± 2.1e+11* |
|  | Seagrass |  | 2.1<sup>\*\*</sup><br>*± 0.6* | 1.3e+09<sup>NS</sup><br>*± 3.0e+08* | 5.8e+13<sup>NS</sup><br>*± 7.8e+12* | 3.7e+11<sup>NS</sup><br>*± 5.6e+10* | 3.8e+12<br>*± 6.1e+11* | 1.5e+11<sup>NS</sup><br>*± 2.8e+10* | 2.4e+11<sup>NS</sup><br>*± 2.8e+10* | 3.6e+10<sup>NS</sup><br>*± 7.7e+09* | 1.5e+12<sup>NS</sup><br>*± 1.7e+11* |
| Ande | Bare | Inner | 3.9<sup>NS</sup><br>*± 1.2* | 8.0e+08<br>*± 7.2e+07* | 2.1e+14<sup>NS</sup><br>*± 2.5e+13* | 2.6e+12<sup>NS</sup><br>*± 2.3e+11* | 2.1e+13<sup>NS</sup><br>*± 1.6e+12* | 7.7e+11<sup>NS</sup><br>*± 9.3e+10* | 1.1e+12<sup>#</sup><br>*± 1.6e+11* | 5.0e+11<sup>\*</sup><br>*± 1.4e+11* | 6.0e+12<sup>NS</sup><br>*± 4.3e+11* |
|  | Seagrass |  | 4.6<sup>NS</sup><br>*± 0.4* | 1.2e+09<sup>#</sup><br>*± 1.8e+08* | 1.7e+14<sup>NS</sup><br>*± 3.5e+13* | 2.0e+12<sup>NS</sup><br>*± 2.1e+11* | 1.7e+13<sup>NS</sup><br>*± 2.3e+12* | 7.9e+11<sup>NS</sup><br>*± 8.5e+10* | 7.7e+11<br>*± 1.1e+11* | 2.1e+11<br>*± 2.8e+10* | 6.1e+12<sup>NS</sup><br>*± 6.7e+11* |
| Font | Bare | Inner | 4.1<sup>NS</sup><br>*± 0.5* | 5.6e+08<br>*± 5.7e+07* | 3.4e+14<sup>NS</sup><br>*± 6.1e+13* | 4.1e+12<sup>#</sup><br>*± 4.4e+11* | 2.9e+13<sup>NS</sup><br>*± 6.6e+12* | 1.6e+12<sup>NS</sup><br>*± 1.6e+11* | 1.0e+12<sup>NS</sup><br>*± 2.1e+11* | 9.5e+11<sup>#</sup><br>*± 8.0e+11* | 9.4e+12<sup>#</sup><br>*± 1.0e+12* |
|  | Seagrass |  | 4.7<sup>NS</sup><br>*± 0.9* | 7.6e+08<sup>#</sup><br>*± 7.7e+07* | 2.7e+14<sup>NS</sup><br>*± 3.6e+13* | 2.9e+12<br>*± 2.8e+11* | 2.5e+13<sup>NS</sup><br>*± 2.8e+12* | 1.5e+12<sup>NS</sup><br>*± 1.9e+11* | 7.7e+11<sup>NS</sup><br>*± 1.1e+11* | 5.7e+11<br>*± 2.7e+11* | 6.8e+12<br>*± 5.3e+11* |
| All | Bare |  | 1.5<br>*± 0.6* | 9.3e+08<br>*± 1.1e+08* | 1.4e+14<sup>NS</sup><br>*± 2.1e+13* | 1.6e+12<sup>NS</sup><br>*± 2.5e+11* | 1.3e+13<sup>NS</sup><br>*± 1.9e+12* | 5.9e+11<sup>NS</sup><br>*± 9.4e+10* | 6.2e+11<sup>NS</sup><br>*± 7.5e+10* | 3.6e+11<sup>NS</sup><br>*± 1.4e+11* | 4.2e+12<sup>NS</sup><br>*± 5.4e+11* |
|  | Seagrass |  | 2.6<br>*± 0.4* | 1.2e+09<sup>\*\*\*</sup><br>*± 1.0e+08* | 1.7e+14<sup>NS</sup><br>*± 1.6e+13* | 1.6e+12<sup>NS</sup><br>*± 1.6e+11* | 1.5e+13<sup>NS</sup><br>*± 1.4e+12* | 6.6e+11<sup>NS</sup><br>*± 8.2e+10* | 6.1e+11<sup>NS</sup><br>*± 5.4e+10* | 2.6 e+11<sup>NS</sup><br>*± 5.1e+10* | 5.1e+12<sup>NS</sup><br>*± 4.3e+11* |
| Location | Inner |  | 4.3<sup>\*\*\*</sup><br>*± 0.4* | 8.4e+08<br>*± 7.2e+07* | 2.5e+14<sup>\*\*\*</sup><br>*± 2.3e+13* | 2.8e+12<sup>\*\*\*</sup><br>*± 2.1e+11* | 2.3e+13<sup>\*\*\*</sup><br>*± 2.0e+12* | 1.2e+12<sup>\*\*\*</sup><br>*± 1.0e+11* | 9.1e+11<sup>\*\*\*</sup><br>*± 7.8e+10* | 5.6e+11<sup>\*\*\*</sup><br>*± 2.1e+11* | 7.1e+12<sup>\*\*\*</sup><br>*± 4.4e+11* |
|  | Outer |  | 0.7<br>*± 0.4* | 1.2e+09\*<br>*± 1.0e+08* | 1.1e+14<br>*± 1.1e+12* | 9.4e+11<br>*± 1.0e+11* | 9.5e+12<br>*± 9.4e+11* | 3.5e+11<br>*± 3.5e+10* | 4.6e+11<br>*± 4.2e+10* | 1.8e+11<br>*± 2.0e+10* | 3.4e+12<br>*± 3.6e+11* |

This spatial organization was mirrored by N_2_O fluxes, which ranged from −3.7 to 8.4 µmol N_2_O m^−2^ d^−1^, with an average value of 2.1 ± 0.4 µmol N_2_O m^−2^ d^−1^ across the bay. Inner sites exhibited approximately six-fold higher emissions than Outer sites (4.3 ± 0.4 vs. 0.7 ± 0.4 µmol N_2_O m^−2^ d^−1^; Wilcoxon test, p = 7.8 × 10^−7^; Figure 3a, Table 1). In contrast, vegetation had no significant overall effect on N_2_O fluxes (Figure 3b, Table 1), although vegetated sediments tended to emit more N_2_O than bare sediments (2.6 ± 0.4 vs. 1.5 ± 0.6 µmol N_2_O m^−2^ d). Significant vegetation effects were detected at only two of the six sites (Wilcoxon test, p < 0.01; Figure 3c, Table 1). Approximately 20% of the measurements showed net N_2_O uptake, occurring predominantly in bare sediments from Outer sites.

**Figure 3.**
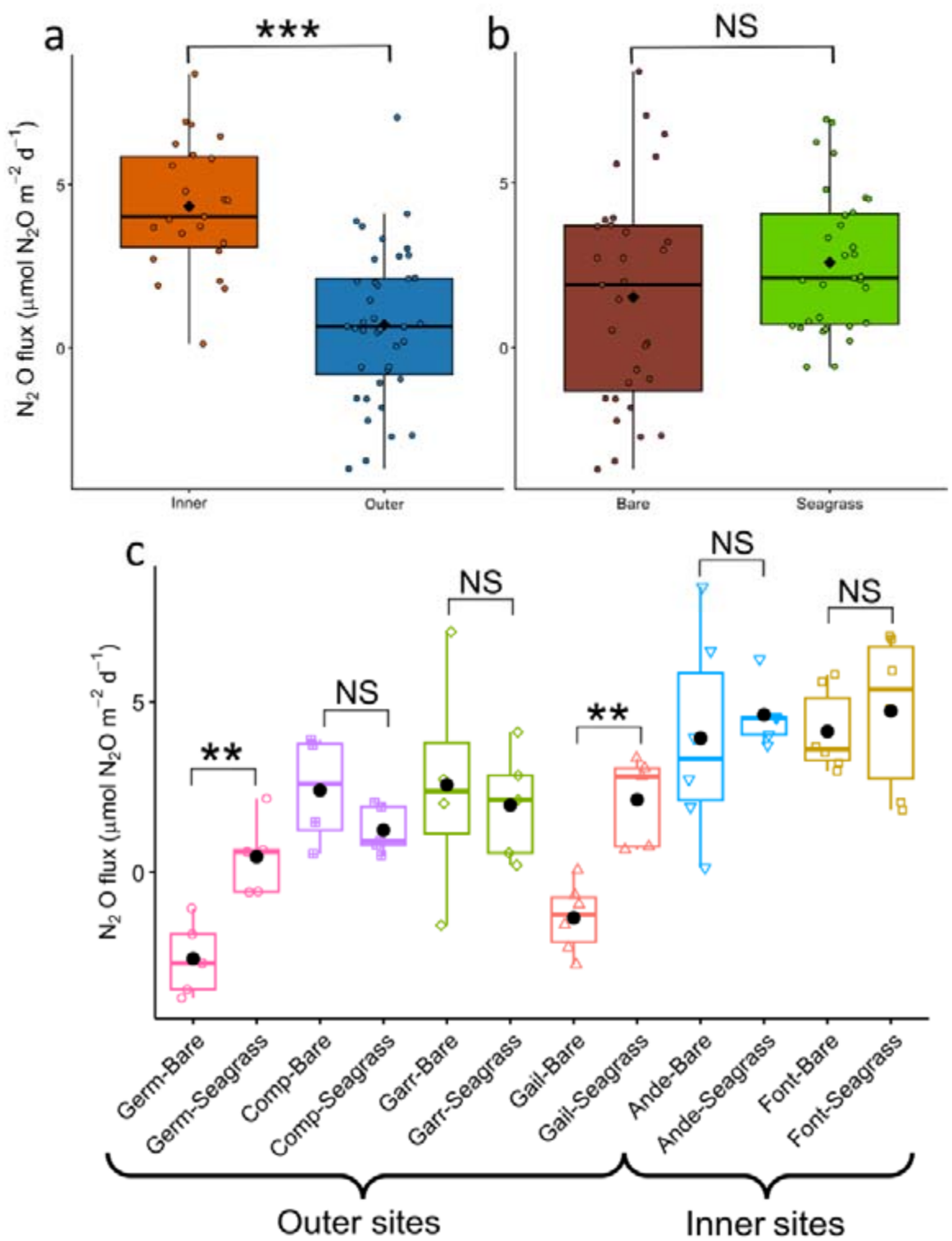
N_2_O fluxes across sites and sediment types in Arcachon Bay. (a) N_2_O fluxes by site location (Inner n = 23, Outer n = 39). (b) by type of sediment (Bare n = 31, Seagrass n = 31). (c) by subsite (n = 6 for Gail-Bare, Font-Bare, and Ande-Bare; n = 5 for Gail-Seagrass, Garr-Seagrass, Ande-Seagrass, Comp-Seagrass, Germ-Bare, and Germ-Seagrass; n = 4 for Garr-Bare and Comp-Bare). In panel (c), sites are sorted by the N_2_O flux of the vegetated sediments.

### 3.2. Structure of N-cycling microbial communities

Of the twelve N-cycling genes investigated, those associated with N fixation were the most abundant (*nifH*; 30.4 ± 0.6%), followed by nitrate reduction (*napA*: NO_3_^−^ → NO_2_^−^; 23.6 ± 0.6%), and nitrite reduction (*nirS*: NO_2_^−^ → NO; 20.5 ± 0.5%), together accounting for approximately 75% of the total relative abundance (Figure S3a). Although their overall distribution remained relatively consistent across sites and vegetation types, PCA analysis of the twelve genes revealed distinct vegetation and spatial patterns (Figure S3b,c). Vegetated sediments (negative side of PC1) were associated with higher relative abundances of *nifH* and *napA*, whereas bare sediments (positive side of PC1) were more closely associated with denitrification genes (PC1 explained ∼39% of the total variance). Spatially, Inner sites (negative side of PC2) were characterized by higher contributions of *nirK, nirS* and *norB*, while Outer sites (the positive side of PC2) were associated with relatively higher abundances of *nrfA, napA, hdh* and *amoA* (PC2 explained ∼23% of the total variance).

Quantification of N_2_O-cycling genes further highlighted spatial differences in microbial functional potential: qPCR showed that *nirK* was more abundant than *nirS*, whereas combining qPCR with targeted metagenomics identified *nirS* as the dominant gene overall (Figure S4). All qPCR-based abundances of N-cycling genes were higher at Inner than at Outer sites, while differences between vegetated and bare sediments were not significant (Table 1). The abundances of the *nir* and *nosZ* genes, as well as their cumulative abundances (*nirKS* and *nosZ12*), were positively correlated with N_2_O fluxes (Figure S5). Similarly to CH_4_-cycling genes in the same studied area^40^, *nir* and *nosZ* gene abundances (qPCR) and gene variant diversity indices were only weakly correlated (Figure S5) and were independently sorted in the general PCA (Figure 2a). The indicates that abundance and diversity describe complementary dimensions of N-cycling microbial communities. The *nir:nosZ* ratios (qPCR-based abundance ratio, ratio_qPCR; richness ratio, Ratio_R; and diversity Ratio, ratio_D) also showed an opposite distribution to diversity indices, with lower ratios being associated with higher community diversity (Figure 2a).

Taxonomic analyses revealed marked differences among functional genes in taxonomic resolution, with *nirK* displaying the highest proportion of classified taxa, whereas approximately 80% of *nosZ1* and *nosZ2* sequence variants remained unassigned at the species level (Table S4). Consequently, subsequent analyses were performed at the genus level. For each of these four genes, the 30 most abundant genera were selected for detailed analysis. Across all four functional genes, genera were consistently more abundant at Inner than at Outer sites and were generally enriched in vegetated sediments at the scale of the bay, with significant vegetation effects detected primarily at 2 sites: Germ and Garr (Figures 4 and S6). Most genera were positively correlated with N_2_O fluxes. The taxonomic composition of N_2_O-cycling communities differed among functional genes (Figures 4 and S6, Supplementary Dataset). The taxa most strongly associated with N_2_O fluxes for *nirS* were predominantly affiliated with Pseudomonadota (formerly Proteobacteria), with additional representatives from Chloroflexota and Planctomycetota. In contrast, *nirK* communities were mainly composed of Thermoproteota and Alphaproteobacteria, with a smaller contribution from Cyanobacteria. The *nosZ1* community was almost exclusively dominated by Pseudomonadota, whereas *nosZ2* communities showed a broader taxonomic composition, including Pseudomonadota, Bacteroidota, Chloroflexota and Balneolota.

**Figure 4.**
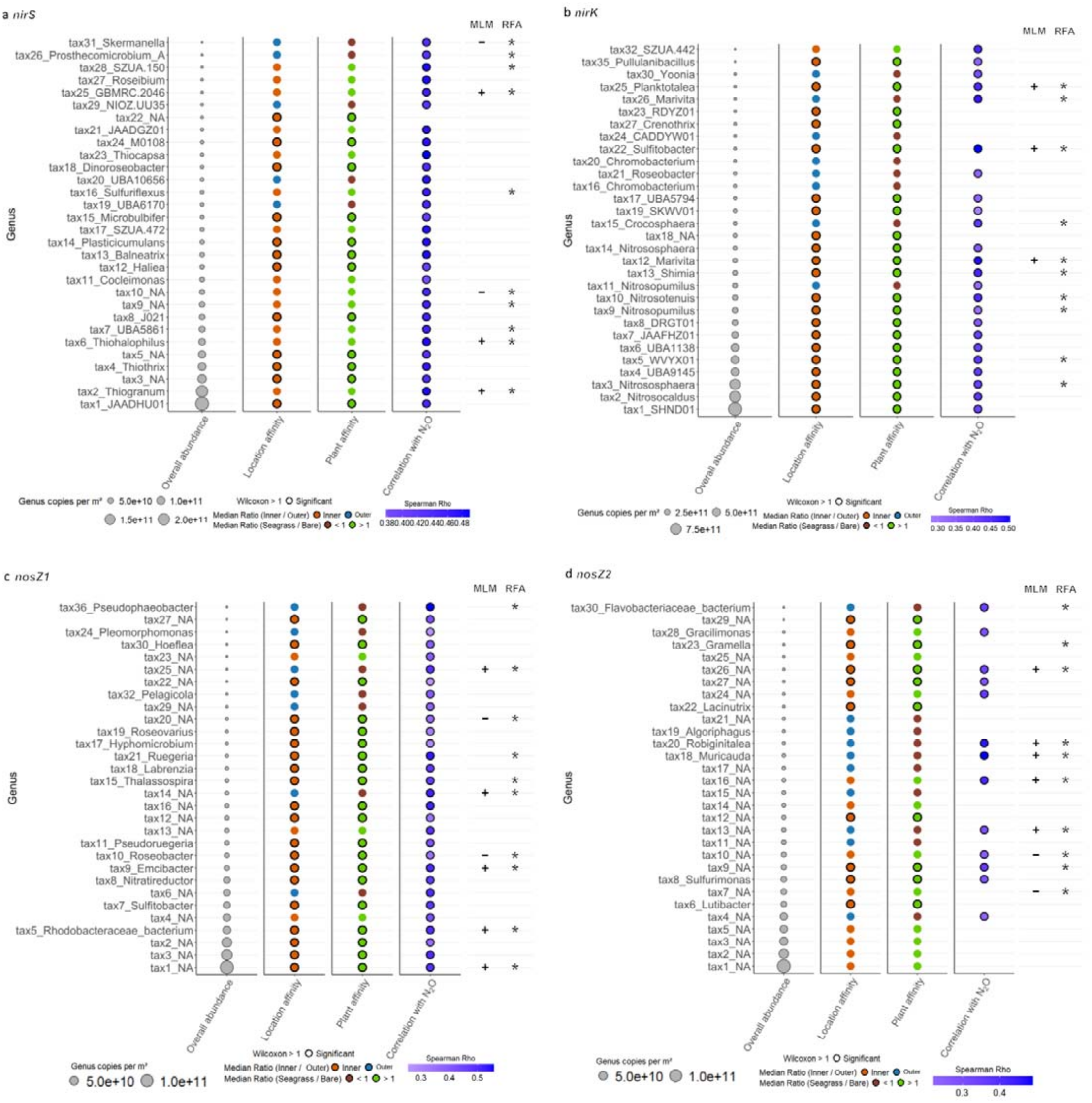
Differential abundance and relationships with N_2_O flux for N_2_O-producing and N_2_O-cycling genera across site location and sediment type in Arcachon Bay. (a–d) Differential abundance and relationships with N_2_O flux for genera carrying the *nirS* (a), *nirK* (b), *nosZ1* (c), and *nosZ2* (d) genes. For each genus, the first column shows mean abundance (gene copies m^−2^) across all samples (n = 71) with point size proportional to abundance. The next two columns show median abundance ratios for Inner/Outer and Seagrass/Bare comparisons. Colours indicate site or sediment-type affinity (orange: Inner/Outer ratio > 1; blue: Inner/Outer ratio < 1; green: Seagrass/Bare ratio > 1; brown: Seagrass/Bare ratio < 1). Significant differences (Wilcoxon test, P < 0.05) are marked by a black circle for Inner or Seagrass affinity. A white star within a blue or brown point indicates Outer or Bare affinity; no such significant differences were observed. The penultimate column shows significant Spearman correlations (ρ) between genus abundance and N_2_O flux, with colour indicating correlation direction and strength (blue: positive). The final column identifies genera selected as important descriptors of N_2_O flux by mixed linear models (MLMs) and random forest analysis (RFA), indicated by positive or negative MLM coefficients and asterisks denoting significance in RFA.

Co-occurrence networks based on sequence variants of *nirK, nirS, nosZ1*, and *nosZ2* showed greater differences in network architecture between Inner and Outer sites than between vegetation types (Figures 5 and S7). Inner networks exhibited a relatively more compact and more connected architecture, with a higher density and clustering coefficient. In contrast, Outer networks displayed greater heterogeneity, modularity and assortativity (Figure 5a,c). Outer networks also contained three times more module hubs than Inner networks (9 vs. 3), including both *nosZ*- and *nir*-associated taxa, while Inner module hubs were exclusively associated with *nir* taxa (Figure 5b). Consistent with these structural differences, robustness analyses showed that the Outer network maintained a higher mean network efficiency when a larger proportion of node were removed, whereas the Inner network displayed a more rapid decline in mean network efficiency, reflecting its greater vulnerability (Figure 5c,d).

**Figure 5.**
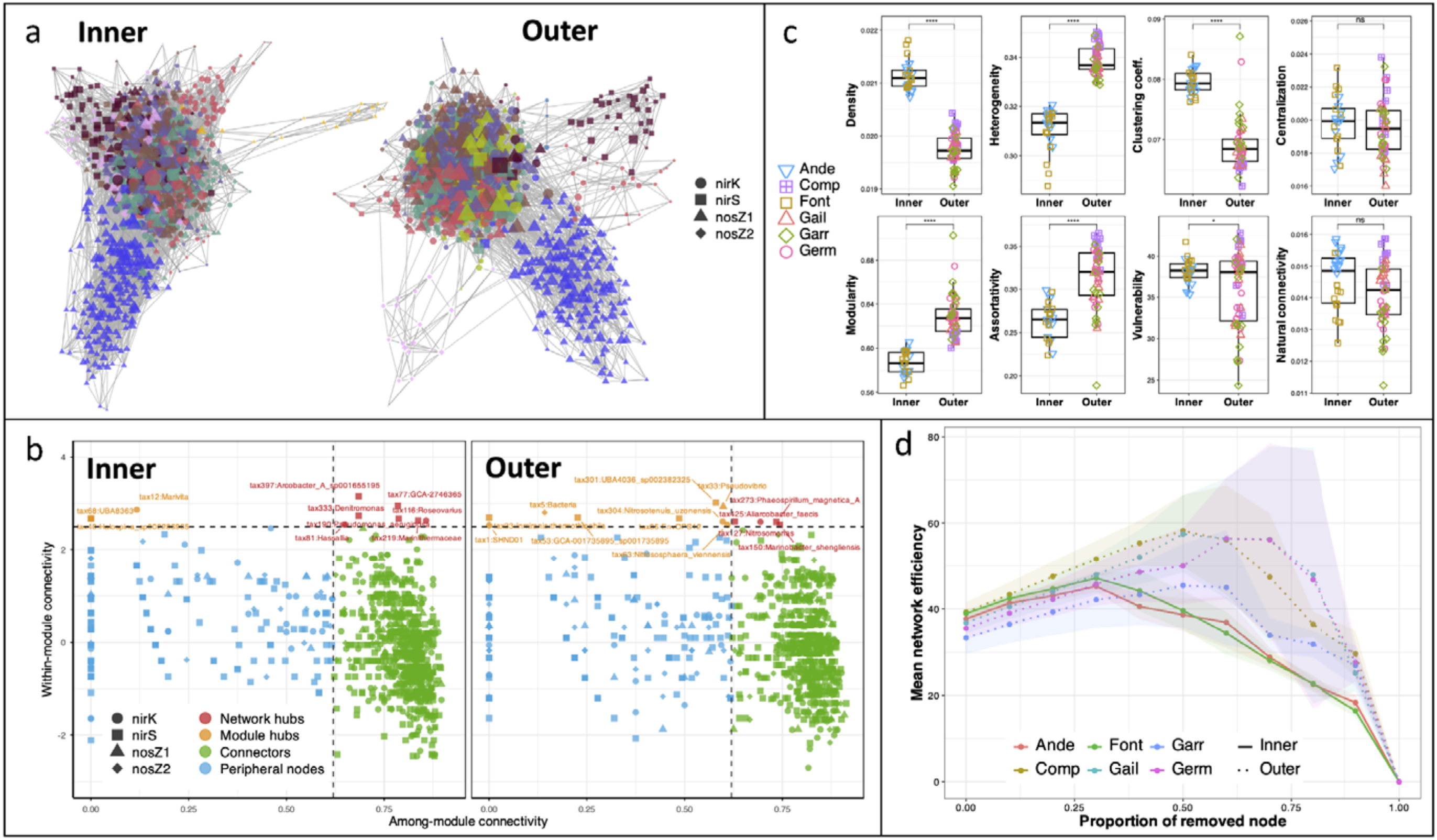
Comparison of microbial co-occurrence network architecture between Inner and Outer sediment communities. (a) Co-occurrence networks constructed for the Inner and Outer sites. Nodes represent microbial taxa, with shapes indicating the associated N_2_O-cycling functional genes (*nirK, nirS, nosZ1*, and *nosZ2*). Node colors indicate the module membership of each taxon. Edges represent significant co-occurrence relationships between taxa. (b) Topological roles of nodes based on within-module connectivity (*Zi*) and among-module connectivity (Pi). Dashed lines indicate the thresholds (*Zi* = 2.5 and *Pi* = 0.62) used to classify nodes as peripheral nodes, connectors, module hubs, or network hubs. Taxa identified as connectors or hubs are annotated. (c) Comparison of network topological properties between Inner and Outer networks, including density, heterogeneity, clustering coefficient, centralization, modularity, assortativity, vulnerability, and natural connectivity. Points represent individual sampling sites, boxplots summarize the distribution of network metrics, and asterisks indicate significant differences between Inner and Outer networks (ns, not significant; ****, *P* < 0.0001). (d) Network robustness assessed by progressive low-degree node removal. Mean network efficiency was calculated from 20 simulations and averaged between 25% and 90% node removal. Colored lines represent the six sampling sites, whereas solid and dotted lines correspond to the Inner and Outer networks, respectively. Shaded areas indicate the variability among simulations.

### 3.3. Environmental and microbial descriptors of N_2_O fluxes

Spatial site variability alone explained 52% of the variance in N_2_O fluxes (Table 2, Model B). This increased to 57% when sites were grouped into Inner and Outer bay (Model D) and reached 59% when vegetation type was incorporated through the Subsite factor (Model A). By contrast, vegetation alone explained only 4% of the variance (Model F), while hierarchical random-effect structures did not substantially improve model performance (Models C and E). Because a large proportion of the variance remained unexplained by spatial categories alone, subsequent analyses explored the contribution of environmental and microbial descriptors. Across both MLMs and RFA, the same descriptors were consistently retained, with network heterogeneity by location emerging as the strongest descriptor of N_2_O fluxes (Tables 2, S2; Figure 6). In the MLM analyses, models were ranked according to AICc, with the best-supported random-effect structure shown in bold, while the remaining models were not significantly different from the best model. The best models explained up to 80% of the N_2_O fluxes variance, and included network property (heterogeneity by location), N-cycling taxa (e.g. *nosZ1 Roseobacter, nirK Planktotalea, nirS Thiohalophilus*), physical (moisture and bathymetry) and chemical (CAR and pH) variables, and richness of *nirKS* (Table 2). When sediment type was included as a random effect, additional taxon-specific descriptors appeared in the best-supported models (model number marked with *). Although the exact combination of descriptors varied with the random-effect structure, several descriptors were consistently retained. The best models including Subsite or Type + Site as random effects (Models 1 and 2) identified network heterogeneity by location, moisture, and *nosZ1* taxon 10 (*Roseobacter*) as negative descriptors of N_2_O fluxes, whereas *nirK* taxon 25 (*Planktotalea*) and pH showed positive relationships. The best models without random effects or including Type, Location, or Type/Location as random effects (Models 5, 11, 12 and 16), consistently retained moisture, bathymetry and *nirKS* richness as negative descriptors, together with positive effects of CAR and pH. When two taxa belonging to the same functional gene were simultaneously included in a model (Models 13, 17 and 18), the regression coefficients increased markedly, suggesting potential collinearity within gene-specific datasets. Regarding microbial descriptors, variables derived from sequencing (*nirKS* richness) were more informative than qPCR variables alone. The best-performing models relied either on descriptors derived from targeted metagenomics alone (diversity metrics and network parameters) or on descriptors combining targeted metagenomics with qPCR-derived gene abundances (taxon-specific absolute abundances), whereas qPCR-derived gene abundances alone were not retained among the best-performing descriptors.

**Table 2.** Summary of the best significant mixed linear models explaining N2O fluxes across different environmental conditions. Models are ranked by 790 Corrected Akaike Information Criterion (AICc), from lowest (best fit) to highest, and are grouped according to fixed or random factors: site, 791 location, sediment type (Bare vs. Seagrass), subsite, Site nested within Type and Location nested within Type. All models include N2O flux as the 792 response variable, with physical, chemical, or microbial variables used as descriptors. Only best models containing all significant (p < 0.05) 793 descriptors within each subset are presented and n = 62. Models marked with an asterisk (*) include taxonomic information. Bold equations 794 indicate the best-performing model within each model structure based on AICc

| Site_ID | Type<br>(Subsite) | Site<br>location | ratio_qPCR<br>copies m<sup>−2</sup> | nirS_R | nirS_D | nirK_R | nirK_D | nosZ1_R | nosZ1_D | nosZ2_R | nosZ2_D | ratio_R | ratio_D |
|---|---|---|---|---|---|---|---|---|---|---|---|---|---|
| Germ | Bare | Outer | 10.5<br>*± 0.6* | 397.7<sup>NS</sup><br>*± 8.1* | 4.4<sup>NS</sup><br>*± 0.1* | 501.2<sup>#</sup><br>*± 20.6* | 5.3<sup>NS</sup><br>*± 0.1* | 1129.7<sup>NS</sup><br>*± 75.0* | 7.0<sup>NS</sup><br>*± 0.1* | 2147.2<sup>NS</sup><br>*± 138.9* | 7.7<sup>NS</sup><br>*± 0.1* | 0.3<sup>NS</sup><br>*± 0.0* | 0.7<sup>NS</sup><br>*± 0.0* |
|  | Seagrass |  | 20.9<sup>\*\*</sup><br>*± 6.9* | 385.3<sup>NS</sup><br>*± 8.1* | 4.4<sup>NS</sup><br>*± 0.0* | 449.5<br>*± 16.5* | 5.2<sup>NS</sup><br>*± 0.1* | 1146.2<sup>NS</sup><br>*± 152.4* | 6.0<sup>NS</sup><br>*± 0.2* | 1939.7<sup>NS</sup><br>*± 267.3* | 7.5<sup>NS</sup><br>*± 0.2* | 0.3<sup>NS</sup><br>*± 0.1* | 0.7<sup>NS</sup><br>*± 0.0* |
| Comp | Bare | Outer | 13.9<sup>NS</sup><br>*± 0.7* | 413.6<sup>NS</sup><br>*± 4.4* | 4.4\*<br>*± 0.0* | 558.0<sup>NS</sup><br>*± 18.7* | 5.3<sup>NS</sup><br>*± 0.1* | 2385.2<sup>\*\*</sup><br>*± 130.8* | 7.8<sup>\*\*</sup><br>*± 0.1* | 2856.4<sup>NS</sup><br>*± 194.4* | 8.0<sup>NS</sup><br>*± 0.1* | 0.2<sup>NS</sup><br>*± 0.0* | 0.6<sup>NS</sup><br>*± 0.0* |
|  | Seagrass |  | 12.5<sup>NS</sup><br>*± 1.2* | 403.0<sup>NS</sup><br>*± 5.2* | 4.3<br>*± 0.0* | 560.2<sup>NS</sup><br>*± 12.0* | 5.4<sup>NS</sup><br>*± 0.1* | 1588.5<br>*± 116.8* | 7.4<br>*± 0.1* | 2813.3<sup>NS</sup><br>*± 200.2* | 7.9<sup>NS</sup><br>*± 0.1* | 0.2<sup>NS</sup><br>*± 0.0* | 0.6<sup>NS</sup><br>*± 0.0* |
| Garr | Bare | Outer | 11.9<sup>NS</sup><br>*± 0.9* | 374.3<sup>NS</sup><br>*± 5.6* | 4.5<sup>#</sup><br>*± 0.0* | 391.5<sup>NS</sup><br>*± 16.9* | 5.2<sup>NS</sup><br>*± 0.0* | 1190.3<sup>NS</sup><br>*± 107.5* | 7.1<sup>NS</sup><br>*± 0.1* | 1560.3<sup>NS</sup><br>*± 126.7* | 7.3<sup>NS</sup><br>*± 0.1* | 0.3<sup>NS</sup><br>*± 0.0* | 0.7<sup>NS</sup><br>*± 0.0* |
|  | Seagrass |  | 13.9<sup>NS</sup><br>*± 2.6* | 386.2<sup>NS</sup><br>*± 12.6* | 4.4<br>*± 0.0* | 453.5<sup>NS</sup><br>*± 46.8* | 5.2<sup>NS</sup><br>*± 0.1* | 1388.3<sup>NS</sup><br>*± 254.8* | 7.1<sup>NS</sup><br>*± 0.3* | 2072.2<sup>NS</sup><br>*± 391.2* | 7.5<sup>NS</sup><br>*± 0.3* | 0.3<sup>NS</sup><br>*± 0.1* | 0.7<sup>NS</sup><br>*± 0.0* |
| Gail | Bare | Outer | 11.8<sup>NS</sup><br>*± 0.8* | 395.3<sup>NS</sup><br>*± 4.9* | 4.3<br>*± 0.0* | 511.3<sup>\*</sup><br>*± 13.8* | 5.3<sup>#</sup><br>*± 0.1* | 2151.3<sup>\*</sup><br>*± 235.6* | 7.6<sup>\*</sup><br>*± 0.1* | 2696.2<sup>\*</sup><br>*± 191.5* | 7.9<sup>\*</sup><br>*± 0.1* | 0.2<br>*± 0.0* | 0.6<br>*± 0.0* |
|  | Seagrass |  | 10.6<sup>NS</sup><br>*± 1.2* | 386.7<sup>NS</sup><br>*± 6.3* | 4.5<sup>\*</sup><br>*± 0.0* | 439.7<br>*± 17.4* | 5.2<br>*± 0.0* | 1290.2<br>*± 166.7* | 7.1<br>*± 0.1* | 2002.8<br>*± 168.0* | 7.6<br>*± 0.1* | 0.3<sup>\*</sup><br>*± 0.0* | 0.7<sup>\*</sup><br>*± 0.0* |
| Ande | Bare | Inner | 12.8<sup>NS</sup><br>*± 1.3* | 397.7<sup>NS</sup><br>*± 8.0* | 4.4<sup>\*</sup><br>*± 0.0* | 559.2<sup>NS</sup><br>*± 20.0* | 5.4<sup>#</sup><br>*± 0.0* | 1170.5<br>*± 129.3* | 7.0<br>*± 0.1* | 2310.5<sup>NS</sup><br>*± 176.7* | 7.7<sup>NS</sup><br>*± 0.1* | 0.3<sup>#</sup><br>*± 0.0* | 0.7<sup>\*</sup><br>*± .0* |
|  | Seagrass |  | 12.3<sup>NS</sup><br>*± 0.9* | 394.2<sup>NS</sup><br>*± 7.4* | 4.3<br>*± 0.0* | 547.0<sup>NS</sup><br>*± 16.6* | 5.3<br>*± 0.1* | 1549.2<sup>#</sup><br>*± 130.7* | 7.3<sup>#</sup><br>*± 0.1* | 2735.3<sup>NS</sup><br>*± 245.1* | 7.9<sup>NS</sup><br>*± 0.1* | 0.2<br>*± 0.0* | 0.6<br>*± 0.0* |
| Font | Bare | Inner | 12.2<sup>NS</sup><br>*± 0.8* | 394.5<sup>NS</sup><br>*± 7.3* | 4.4<sup>NS</sup><br>*± 0.0* | 524.7<sup>NS</sup><br>*± 27.4* | 5.3<sup>NS</sup><br>*± 0.1* | 1658.0<sup>NS</sup><br>*± 168.9* | 7.4<sup>NS</sup><br>*± 0.1* | 2281.2<br>*± 153.4* | 7.7<br>*± 0.1* | 0.2<sup>\*\*</sup><br>*± 0.0* | 0.6<sup>#</sup><br>*± 0.0* |
|  | Seagrass |  | 12.6<sup>NS</sup><br>*± 0.5* | 390.5<sup>NS</sup><br>*± 9.5* | 4.3<sup>NS</sup><br>*± 0.0* | 549.5<sup>NS</sup><br>*± 18.9* | 5.4<sup>NS</sup><br>*± 0.0* | 1918.3<sup>NS</sup><br>*± 83.1* | 7.6<sup>NS</sup><br>*± 0.0* | 3134.2<sup>\*\*</sup><br>*± 151.3* | 8.0<sup>\*\*</sup><br>*± 0.1* | 0.2<br>*± 0.0* | 0.6<br>*± 0.0* |
| All | Bare |  | 12.1<sup>NS</sup><br>*± 0.4* | 395.0<sup>NS</sup><br>*± 3.2* | 4.4<sup>\*\*\*</sup><br>*± 0.0* | 506.2<sup>NS</sup><br>*± 12.3* | 5.3<sup>NS</sup><br>*± 0.0* | 1592.1<sup>NS</sup><br>*± 101.4* | 7.3<sup>NS</sup><br>*± 0.1* | 2293.0<sup>NS</sup><br>*± 93.8* | 7.7<sup>NS</sup><br>*± 0.0* | 0.7<sup>NS</sup><br>*± 0.0* | 0.7<sup>NS</sup><br>*± 0.0* |
|  | Seagrass |  | 13.8<sup>NS</sup><br>*± 1.3* | 391.0<sup>NS</sup><br>*± 3.4* | 4.3<br>*± 0.0* | 500.0<sup>NS</sup><br>*± 12.8* | 5.3<sup>NS</sup><br>*± 0.0* | 1480.1<sup>NS</sup><br>*± 73.4* | 7.2<sup>NS</sup><br>*± 0.1* | 2449.6<sup>NS</sup><br>*± 122.7* | 7.7<sup>NS</sup><br>*± 0.1* | 0.3<sup>NS</sup><br>*± 0.0* | 0.7<sup>NS</sup><br>*± 0.0* |
| Location | Inner |  | 12.5<sup>NS</sup><br>*± 0.4* | 394.2<sup>NS</sup><br>*± 3.8* | 4.3<br>*± 0.0* | 545.1<sup>\*\*\*</sup><br>*± 10.2* | 5.4<sup>\*</sup><br>*± 0.0* | 1574.0<sup>NS</sup><br>*± 83.1* | 7.3<sup>NS</sup><br>*± 0.1* | 2615.3<sup>\*</sup><br>*± 113.1* | 7.8<sup>\*</sup><br>*± 0.0* | 0.2<br>*± 0.0* | 0.6<br>*± 0.0* |
|  | Outer |  | 13.2<sup>NS</sup><br>*± 1.0* | 392.3<sup>NS</sup><br>*± 2.9* | 4.4<sup>\*</sup><br>*± 0.0* | 481.5<br>*± 11.1* | 5.3<br>*± 0.0* | 1515.6<sup>NS</sup><br>*± 84.4* | 7.2<sup>NS</sup><br>*± 0.1* | 2278.3<br>*± 97.8* | 7.7<br>*± 0.1* | 0.3<sup>\*</sup><br>*± 0.0* | 0.7<sup>\*</sup><br>*± 0.0* |

**Figure 6.**
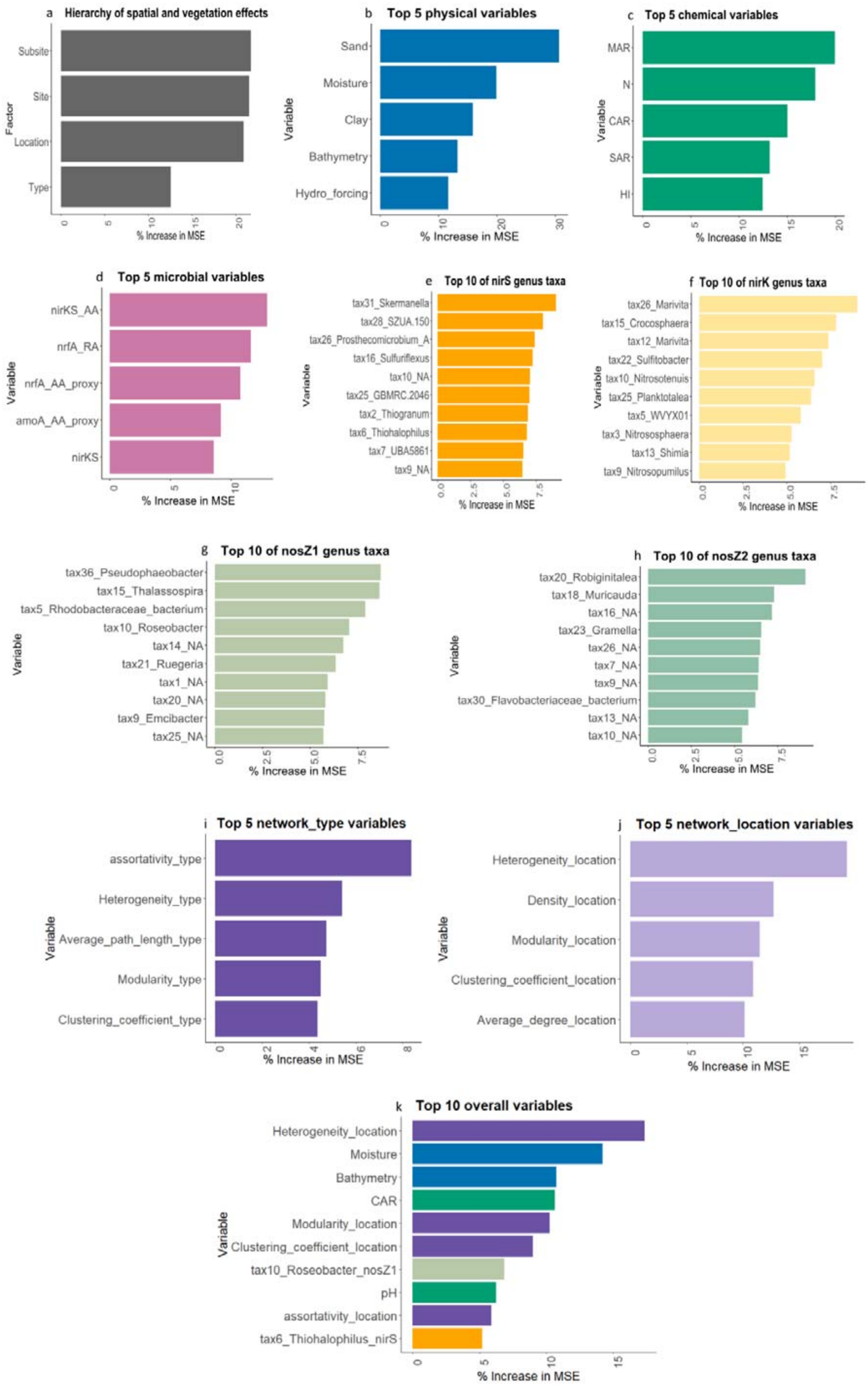
Variable importance in random forest analysis identifying key descriptors of N_2_O fluxes. (a) Hierarchy of spatial and vegetation effects (site, location, sediment type, and subsite). (b-c) Top 5 descriptors within each environmental category: (b) physical, (c) chemical variables, n = 62. (d) Top 5 microbial descriptors. (e-h) Top 10 taxonomic descriptors at the genus level for *nirS, nirK, nosZ1* and *nosZ2* genes, respectively. (i-j) Top 10 microbial network by type and by location, respectively. (k) Overall top 10 descriptors of N_2_O flux across all variables.

## 4. DISCUSSION

N_2_O fluxes measured in Arcachon Bay ranged from net consumption to moderate emissions, consistent with the range reported for seagrass ecosystems^5,10,28^. Low or negative N_2_O fluxes are frequently observed in seagrass sediments where efficient N-cycling can favour complete denitrification, although substantial variability occurs depending on nutrient and OM availability^4,10,28,41^. Approximately 20% of our measurements showed net N_2_O consumption, mainly in bare sediments, suggesting that microbial N_2_O reduction to N_2_ may contribute to these net sinks^42,43^. However, comparisons with previous studies remain challenging, because coastal assessments often rely on water–air fluxes, whereas our measurements captured sediment–air exchanges during low tide. Moreover, all these measurements represent short-term snapshots and do not capture diel, tidal or seasonal variability, all of which can influence N_2_O fluxes through shifts in oxygen availability, microbial activity and seagrass growth^10,44–47^. Therefore, the reported daytime low-tide fluxes measured during the seasonal biomass peak of *Z. noltii* should be interpreted as snapshots of potentially elevated emission periods rather than complete 24-hour cycles. Longer-term measurements integrating tidal and seasonal dynamics will be required to better constrain coastal N_2_O budgets.

Despite this temporal limitation, spatial variability strongly structured N_2_O dynamics in Arcachon Bay. Site position was a stronger descriptor of N_2_O fluxes than vegetation, with Inner sites exhibiting approximately six-fold higher emissions than Outer sites (Table 1). This spatial separation was also captured by PCA (Figure 2), which integrated differences in hydrodynamic exposure, sediment characteristics, C availability, microbial functional potential and community structure. Thus, in contrast to CH_4_ dynamics in the same system, for which vegetation was a dominant descriptor^40^, N_2_O emissions were primarily associated with the environmental context in which seagrass occurred. Higher emissions at Inner sites were linked to greater labile C availability and distinct sedimentary conditions. These sites exhibited higher labile C and C burial, together with greater bacterial abundance and higher abundances of N-cycling in shallower, drier and coarser sediments. Similar patterns have been reported in aquatic sediments^10,18,48^, where reduced hydrodynamic flushing in semi-enclosed environments promotes OM retention and microbial N transformations while sandy sediments may enhance oxygen penetration and generate dynamic oxic-anoxic transitions, favouring coupled nitrification–denitrification and transient N_2_O accumulation^49,50^.

Although seagrass sediments occasionally showed higher N_2_O emissions, the effects of vegetation were strongly site-dependent. Seagrass modifies oxygen availability, organic inputs and rhizosphere redox conditions, but these effects depend on the surrounding environmental conditions^51^. In Outer sites, more C is more limited, the role of vegetation seems to be enhanced. Our results therefore suggest that seagrass acts primarily as a local habitat modifier, whereas sediment properties, C availability and hydrodynamic exposure provide the dominant environmental framework controlling N_2_O emissions.

The spatial differences observed in N_2_O fluxes were accompanied by marked differences in the abundance and composition of N-cycling microbial communities. Both targeted metagenomics and qPCR revealed clear spatial structuring of N-cycling microbial communities, with Inner sites generally exhibiting higher abundances of N-cycling genes (Figure 2, Table 1). N_2_O fluxes increased with labile C and C burial, whereas higher sediment N content was associated with lower fluxes. Together with the strong positive relationships between N-cycling gene abundances, bacterial abundance and labile C, these patterns suggest that microbial N transformations were primarily shaped by C availability rather than N limitation^52^. This interpretation is consistent with the relatively high N inputs received by Arcachon Bay from its catchment, which may maintain sufficient N availability within the system^53^. C availability can control microbial N transformations in aquatic sediments by regulating the supply of electron donors required for nitrate reduction^48^.

The contrasting distribution of denitrification- and DNRA-associated genes further suggests differences in the N fate between site groups. Denitrification genes were generally enriched at Inner sites, whereas *nrfA*, used here as a molecular proxy for DNRA, was more abundant at Outer sites (Figure S3b). The positive relationship between *nrfA* and sediment N content, combined with its marginal negative association with N_2_O fluxes is consistent with a greater potential for N retention through DNRA at Outer sites. Conversely, the higher abundance of denitrification genes at Inner sites suggests a greater potential for gaseous N loss. However, gene abundance reflects functional potential rather than activity and cannot directly demonstrate differences in pathway rates.

Differences in C availability, sediment texture and hydrodynamic forcing likely contributed jointly to this microbial structuring. Inner sites accumulated more carbon under reduced hydrodynamic disturbance, providing electron donors that may stimulate heterotrophic denitrification, whereas Outer sites experienced stronger sediment mixing and reduced OM retention. These environmental contrasts likely generated different redox conditions and microbial niches, shaping both the composition and functional potential of N-cycling communities^48,54^. Higher sediment accumulation rates at Inner sites may favour fast-growing denitrifiers over slow-growing DNRA^55^.

The influence of vegetation on N-cycling microbial communities were more context-dependent. Although seagrass sediments were enriched in several N-cycling taxa across the bay, vegetation effects on community composition varied markedly among sampling sites rather than following a consistent spatial pattern (Figures 4 and S6). Seagrass-derived organic compounds, radial oxygen loss and rhizosphere redox gradients can create distinct microbial niches^56,57^, but their influence depends on the surrounding sedimentary environment. Consequently, seagrass primarily promotes specific microbial taxa locally, whereas broader sedimentary and hydrodynamic conditions govern the overall organization of N-cycling communities.

Importantly, higher N-cycling gene abundance at Inner sites did not directly explain their greater N_2_O emissions. Efficient coupling between upstream denitrification steps and N_2_O reduction can prevent N_2_O accumulation despite high functional potential, highlighting the need to consider microbial diversity and organization beyond gene abundance alone.

Differences in N_2_O fluxes between Inner and Outer sites were associated not only with variation in N-cycling gene abundance, but also with differences in microbial community composition and organization. Among all the physicochemical, functional and microbial variables considered in the combined models, network heterogeneity emerged as the strongest descriptor of N_2_O flux variation, highlighting the relevance of microbial community organization as an additional layer of ecosystem functioning. Co-occurrence networks displayed distinct topological characteristics between site groups, with network heterogeneity being negatively associated with N_2_O fluxes. Although co-occurrence patterns do not demonstrate direct ecological interactions, microbial network architecture captures emergent properties of community organization beyond taxonomic composition alone^24,25^. In line with our findings, recent studies has suggested that microbial interaction networks may provide additional insights into N_2_O emissions by integrating higher-order properties of microbial communities beyond taxonomic or functional composition alone^23^. Lower network heterogeneity at Inner sites may reflect more stable and depositional sedimentary conditions, characterized by reduced hydrodynamic disturbance and greater OM accumulation, resulting in more homogeneous microbial assemblages. Conversely, stronger tidal flushing and sediment resuspension at Outer sites likely generate a wider diversity of microhabitats, favouring more heterogeneous and compartmentalized microbial networks^58^. Such organization may enhance the persistence of microbial functions by maintaining multiple taxa capable of performing similar ecological roles, thereby increasing functional redundancy and potentially supporting more efficient N transformations under variable environmental conditions.

Taxonomic composition further supported differences in microbial community organization. N-cycling communities were dominated by Proteobacteria, particularly Gammaproteobacteria, together with sulfur-associated groups commonly found in organic-rich and redox-variable marine sediments. Several sulfur-oxidizing taxa can couple sulfur oxidation to nitrate reduction processes, including denitrification, highlighting the importance of coupled biogeochemical pathways in these environments^59–63^.

Targeted metagenomics revealed a high prevalence of *nirS*-associated taxa across sites, consistent with their frequent dominance in aquatic environments^64,65^. The prevalence of *nirS* containing microorganisms may reflect their metabolic flexibility, including the ability to couple denitrification with alternative redox processes such as sulfur, iron or arsenate reduction, providing an advantage in fluctuating tidal sediments^66^. Beyond gene abundance, higher diversity within *nir*-associated communities was associated with lower N_2_O fluxes, suggesting that greater functional redundancy among N-cycling microorganisms may enhance the stability and efficiency of N_2_O reduction processes^67^. The positive relationship between *nosZ* abundance and N_2_O fluxes likely reflects the enrichment of N_2_O reducers in environments with greater substrate availability rather than a direct stimulation of emissions. In addition, contrasting nirK/nirS patterns between qPCR and targeted sequencing highlight the complementarity of approaches based on quantitative gene abundance and community diversity, given methodological differences, primer specificity and sequence capture^38,68^.

Overall, our results indicate that N_2_O emissions originate from multiple levels of microbial organization. While N-cycling gene abundance reflects functional potential, network architecture, taxonomic composition and diversity provide complementary information on how microbial communities regulate the balance between N_2_O production and reduction. However, DNA-based approaches capture functional potential rather than activity, highlighting the need for metatranscriptomic approaches to identify the active microbial processes driving N_2_O dynamics.

In conclusion, our study demonstrates that spatial environmental heterogeneity, rather than seagrass presence alone, is the primary descriptor of N_2_O flux variability across Arcachon Bay. Hydrodynamic exposure, sediment characteristics and C availability shape microbial N-cycling communities and their network architecture, highlighting the importance of environmental context in regulating N_2_O dynamics. Beyond functional gene abundance, microbial community organization and diversity provided complementary insights into N_2_O regulation, with greater network heterogeneity and *nir* diversity associated with lower emissions. These findings highlight the importance of integrating microbial community characteristics with environmental drivers to better understand coastal N_2_O dynamics. Future research combining high-resolution N_2_O flux measurements and microbial community analyses will improve our understanding of the mechanisms controlling N_2_O emissions and assess the climate mitigation potential of blue C ecosystems more accurately.

## Acknowledgements

We thank Ifremer, PNMBA, and Pierrick Fribourg for providing nautical resources. We are also grateful to the two interns, Rémi Dugué and Margot Ahr, for their valuable assistance during the fieldwork and laboratory analyses. Finally, we thank the AME (Activités Microbiennes dans l’Environnement) platform of the Laboratory of Microbial Ecology (LEM, Lyon, France) for its support with microbial activity measurements.

## Conflicts of interest

None declared.

## Funding

This work was supported by the Green Deal project REST-COAST, funded by the European Union’s Horizon 2020 research and innovation programme (Grant Agreement 101037097). Additional support was provided by the Graduate School H2O’Lyon (ANR-17-EURE-0018) of Université de Lyon (UdL), within the program *“Investissements d’Avenir”* operated by the French National Research Agency (ANR). HS and DP was funded by the Research Council of Finland in NitroBiome (nos. 342362, 342362, 361980), and the Atmosphere and Climate Competence Center (ACCC) (nos. 337550, 357905, 359343).

## Data and code availability

Radionclide data used to calculate the SAR and MAR are archived on the SEANOE platform [https://doi.org/10.17882/99990]. The Excel file and R code, which include the other variables used in this study and those for figure generation, are archived on Zenodo at [https://doi.org/10.5281/zenodo.23477486]. The Zenodo dataset is currently under embargo but is accessible to reviewers via the following private link during the peer-review process: Private reviewer link. The dataset will be made publicly available upon publication of the associated article or at the end of the embargo period, whichever occurs first. The Excel file contains data collected from six sites within Arcachon Bay, and the R code includes statistical analyses (e.g. Wilcoxon tests, mixed linear models, and random forest analyses) as well as scripts for figure generation. Additionally, raw targeted metagenomic sequences generated in this study have been deposited in the NCBI Sequence Read Archive (SRA) under BioProject accession number PRJNA1335804.

---

| **Model** | **Formula: Fixed part** | **Random part** | **AICc (GLS)** | **Fixed R<sup>2</sup>** | **Global R<sup>2</sup>** |
|---|---|---|---|---|---|
| A |  | ~1\|Subsite | 154 | 0 | 0.59 |
| B |  | ~1\|Site | 155 | 0 | 0.52 |
| C |  | ~1\|Type/Site | 156 | 0 | 0.59 |
| D |  | ~1\|Location | 158 | 0 | 0.57 |
| E |  | ~1\|Type/Location | 161 | 0 | 0.50 |
| F |  | ~1\|Type | 183 | 0 | 0.04 |
| **1\*** | **N<sub>2</sub>O_flux = - 1.41 x Heterogeneity_location - 1.15 X tax10_Roseobacter_nosZ1 - 1.13 X sqrt(Moisture) + 0.94 x tax25_Planktotalea_nirK + 0.79 x pH** | **~1\|Subsite** | **258** | **0.53** | **0.77** |
| **2\*** | **N<sub>2</sub>O_flux = - 1.38 x Heterogeneity_location - 1.32 x sqrt(Moisture) - 1.27 x tax10_Roseobacter_nosZ1 + 0.90 x tax25_Planktotalea_nirK + 0.74 x pH** | **~1\|Type/Site** | **259** | **0.50** | **0.80** |
| 3* | N<sub>2</sub>O_flux = - 1.41 x Heterogeneity_location - 1.24 x sqrt(Moisture) - 1.01 x tax10_Roseobacter_nosZ1 + 0.72 x pH + 0.71 x tax25_NA_nosZ1 | ~1\|Type/Site | 259 | 0.49 | 0.80 |
| 4* | N<sub>2</sub>O_flux = - 1.47 x Heterogeneity_location - 1.00 X sqrt(Moisture) - 0.84 x tax10_Roseobacter_nosZ1 + 0.77 x pH + 0.71 x tax25_NA_nosZ1 | ~1\|Subsite | 259 | 0.53 | 0.76 |
| **5** | **N<sub>2</sub>O_flux = - 1.35 x sqrt(Moisture) - 1.20 x Bathymetry - 0.95 x nirKS_R + 0.88 x CAR + 0.79 x pH** |  | **253(259)** | **0.62** | **0.62** |
| 6 | N<sub>2</sub>O_flux = - 1.25 x Heterogeneity_location - 0.94 X sqrt(Moisture) + 0.82 x pH - 0.71 x nirKS_R + 0.69 x tax25_NA_nosZ1 | ~1\|Subsite | 260 | 0.55 | 0.75 |
| **7** | **N<sub>2</sub>O_flux = - 1.30 X sqrt(Moisture) - 1.22 x Heterogeneity_location + 0.88 x pH + 0.85 x CAR - 0.82 x nirKS_R** | **~1\|Site** | **260** | **0.61** | **0.73** |
| 8* | N<sub>2</sub>O_flux = - 1.75 x Heterogeneity_location - 0.95 x tax5_Rhodobacteraceae_bacterium_nosZ1 + 0.90 x tax25_NA_nosZ1 - 0.85 X sqrt(Moisture) + 0.77 x pH | ~1\|Subsite | 260 | 0.53 | 0.72 |
| 9* | N<sub>2</sub>O_flux = - 1.77 x Heterogeneity_location - 1.01 x tax6_Thiohalophilus_nirS - 0.99 X sqrt(Moisture) + 0.78 x tax25_NA_nosZ1 + 0.72 x pH | ~1\|Subsite | 260 | 0.53 | 0.71 |
| 10* | N<sub>2</sub>O_flux = - 1.37 X sqrt(Moisture) - 1.28 x nirKS_R + 0.87 x CAR + 0.77 x pH + 0.75 x tax25_NA_nosZ1 | ~1\|Site | 261 | 0.49 | 0.70 |
| **11** | **N<sub>2</sub>O_flux = - 1.35 x sqrt(Moisture) - 1.20 x Bathymetry - 0.95 x nirKS_R + 0.88 x CAR + 0.79 x pH** | **~1\|Type** | **262** | **0.64** | **0.64** |
| **12** | **N<sub>2</sub>O_flux = - 1.35 x sqrt(Moisture) - 1.20 x Bathymetry - 0.95 x nirKS_R + 0.88 x CAR + 0.79 x pH** | **~1\|Location** | **262** | **0.64** | **0.64** |
| 13* | N<sub>2</sub>O_flux = 3.62 x tax6_Thiohalophilus_nirS - 3.59 x tax10_NA_nirS - 1.63 x sqrt(Moisture) - 1.15 x Heterogeneity_location - 1.08 x tax10_Roseobacter_nosZ1 | ~1\|Type | 263 | 0.45 | 0.74 |
| 14* | N<sub>2</sub>O_flux = - 1.81 x Heterogeneity_location - 1.35 x tax5_Rhodobacteraceae_bacterium_nosZ1 - 1.28 x sqrt(Moisture) + 0.76 x pH + 0.91 x tax25_NA_nosZ1 | ~1\|Type | 264 | 0.53 | 0.68 |
| 15* | N<sub>2</sub>O_flux = - 1.72 x Heterogeneity_location - 1.40 x sqrt(Moisture) - 1.19 x tax10_NA_nirS + 0.62 x pH + 0.58 x tax25_NA_nosZ1 | ~1\|Type | 264 | 0.55 | 0.67 |
| **16** | **N<sub>2</sub>O_flux = - 1.35 x sqrt(Moisture) - 1.20 x Bathymetry - 0.95 x nirKS_R + 0.88 x CAR + 0.79 x pH** | **~1\|Type/Location** | **265** | **0.64** | **0.64** |
| 17* | N<sub>2</sub>O_flux = 4.14 x tax6_Thiohalophilus_nirS - 3.59 x tax10_NA_nirS - 1.58 x tax10_Roseobacter_nosZ1 - 1.37 x sqrt(Moisture) + 0.79 x pH | ~1\|Type/Location | 266 | 0.45 | 0.74 |
| 18* | N<sub>2</sub>O_flux = 3.62 x tax6_Thiohalophilus_nirS - 3.59 x tax10_NA_nirS - 1.63 x sqrt(Moisture) - 1.15 x Heterogeneity_location - 1.08 x tax10_Roseobacter_nosZ1 | ~1\|Type/Location | 266 | 0.25 | 0.71 |
| 19* | N<sub>2</sub>O_flux = - 1.81 x Heterogeneity_location - 1.35 x tax5_Rhodobacteraceae_bacterium_nosZ1 - 1.28 x sqrt(Moisture) + 0.91 x tax25_NA_nosZ1 + 0.76 x tax10_NA_nirS | ~1\|Type/Location | 266 | 0.53 | 0.68 |
| 20* | N<sub>2</sub>O_flux = - 1.72 x Heterogeneity_location - 1.40 x sqrt(Moisture) - 1.19 x tax10_NA_nirS + 0.62 x pH + 0.58 x tax25_NA_nosZ1 | ~1\|Type/Location | 267 | 0.55 | 0.67 |

## Notes

https://doi.org/10.5281/zenodo.23477486

https://doi.org/10.17882/99990

### Competing Interest Statement

The authors have declared no competing interest.

