## Supplementary M&M and tables/figures for "Environmental heterogeneity and microbial community architecture shape N_2_O fluxes in intertidal coastal sediments"

**Supplementary text**

**Materials and Methods**

**Characterisation of the sediments and sites**

**Plant cover**

Seagrass (*Zostera noltii*) coverage was quantified following the European Water Framework Directive protocol^1^. At each study point, a 30 × 30 cm quadrat was placed, photographed, and the percentage of seabed covered by seagrass was independently estimated by three researchers. The final coverage value for each quadrat was calculated as the average of these three estimates.

**Hydrodynamic forcing**

Hydrodynamic forcing was assessed using a dimensionless parameter, which quantifies the total hydrodynamic energy during a tidal cycle (spring or neap tide) that exceeds a critical velocity threshold for sediment erosion or leaf uprooting. This parameter was calculated following sediment transport model formulations^2^, and applies to both vegetated and unvegetated areas. Further methodological details are available in Dolivet-Maréchal et al.^3^.

**Analyses on the top 10 cm of mixed sediment cores**

**Abundance of bacterial 16S rRNA and functional N_2_O-cycling genes**

DNA was extracted from sediment samples using the NucleoSpin® Soil Kit (Macherey-Nagel GmbH & Co, Düren, Germany)^4^ following the manufacturer’s protocol. DNA concentrations were quantified using a Qubit® 2.0 fluorometer and the Quant-iT™ dsDNA Broad Range (BR) Assay Kit (Invitrogen, France). The abundance of the bacterial community was estimated by absolute quantification of 16S rRNA gene copy number via quantitative real-time PCR (qPCR), using iTaq™ Universal SYBR® Green Supermix (Bio-Rad) on a CFX Connect Real-Time PCR Detection System (Bio-Rad). The abundance of key genes involved in N_2_O production, i.e. *nirS* and *nirK*, and N_2_O consumption, i.e. *nosZ1* and *nosZ2* were also quantified. Detailed information on thermal conditions and primer sets is provided in Table S1. Melting curve analysis was conducted to verify PCR product specificity after amplification. Amplification showed a linear response (R² > 0.99) with no detectable PCR inhibitors, which were systematically checked.

**Targeted metagenomic bioinformatic pipeline**

Gene-specific probes were designed from clustered reference sequences at predefined similarity thresholds^5^. DNA sequencing was performed using the Illumina platform (Arbor Biosciences, Ann Arbor, MI, USA), producing approximately five million reads per sample. Raw sequences were processed using in-house bioinformatic pipelines on the CSC Puhti supercomputer (Espoo, Finland). Gene sequences were identified using^6^ HMMER v3.3.2 via the nhmmer tool with a threshold of E < 0.0001. Multiple sequence alignments were conducted with MAFFT^7^, and phylogenetic placements were inferred with RAxML (version 8.2.12)^8^.

Relative abundances of the 12 targeted N-cycling genes were calculated as the proportion of reads affiliated with each gene relative to the total reads detected for these 12 genes in each sample. For *nirS*, *nirK*, *nosZ1* and *nosZ2*, these relative abundances were multiplied by qPCR-derived gene copy numbers to estimate absolute abundances (*gene* _AA; e.g. *nirS*_AA). Because qPCR data were not available for *amoA* and *nrfA*, absolute abundance proxies (*amoA*_proxy_AA and *nrfA*_proxy_AA) were estimated using a cross-product approach based on the four quantified genes.

Relative taxonomic abundances were calculated from sequencing reads and converted into absolute abundances using gene-specific qPCR estimates. For each functional gene, the 30 genera with the highest absolute abundances across all samples were retained for subsequent analyses. Community composition was visualized at the genus level using ggplot2 (v 4.0.1) in R.

Alpha diversity metrics were calculated separately for each sample to assess the diversity of *nirS*, *nirK*, *nosZ1* and *nosZ2* gene sequences. Richness (number of unique placements) and Shannon diversity index were computed using the nplace count per phylogenetic placement as a proxy for abundance. These calculations were performed using the R package *vegan* (v1.44.0)^9^, applying the specnumber() and diversity() functions.

**Co-occurrence networks**

Co-occurrence networks were constructed to explore potential associations among microbial taxa and to compare network architecture between sampling locations (Inner- vs. Outer-bay) and sample type (Bare- vs. Seagrass sediments). Separate networks were inferred for condition after retaining microbial taxa detected in at least 50% of the corresponding samples. Networks were inferred using the spiec-easi R package (version 1.99.0)^10^, following the methodology previously described Schwob et al.^11^. Sample-specific subnetworks were extracted from each condition-level network using the subset_network() function in *meconetcomp* R package (version 0.7.1), retaining only taxa detected in each sample^12^. Topological properties were calculated for each sample-specific network using *meconetcomp*. Natural connectivity was calculated with *pulsar* R package (version 0.3.13)^13^. Degree assortativity was calculated with *igraph* R package (version 2.2.2)^14^. Differences in network properties between Inner- and Outer-bay samples and between bare and vegetated sediments were assessed using approximate permutation tests implemented in the coin R package, with 9,999 permutations^15^. Node topological roles were determined from within-module connectivity (*Zi*) and among-module participation (*Pi*). Nodes were classified as peripheral nodes (*Zi* ≤ 2.5 and *Pi* ≤ 0.62), connectors (*Zi* ≤ 2.5 and *Pi* > 0.62), module hubs (*Zi* > 2.5 and *Pi* ≤ 0.62), or network hubs (*Zi* > 2.5 and *Pi* > 0.62)^16,17^. Network robustness was evaluated using 20 simulations of progressive low-degree node removal with the robustness() function in *meconetcomp*. Efficiency values were averaged between 25% and 90% node removal, providing an integrated measure of network connectivity during the loss of the least-connected taxa.

**Random Forest analysis and mixed linear models**

RFA was applied separately to each predefined subset of descriptors using the *randomForest* R package. Model tuning was performed using the tuneRF() function by testing different *mtry* values, corresponding to the number of variables randomly sampled at each split, across 1000 trees. At each iteration, *mtry* was adjusted by a factor of 1.5, and the search continued only when the relative improvement in out-of-bag (OOB) error exceeded 1%. This procedure allowed the identification of the optimal *mtry* value while limiting overfitting. Variable importance was assessed using the mean increase in mean squared error (%IncMSE). Variables showing the highest importance scores within each subset were retained as key descriptors and compared with those identified by MLM analyses. This combined approach allowed cross-validation of the main descriptors of N₂O fluxes and ensured that retained descriptors were supported by both inferential and machine-learning approaches.

**Moisture**

Moisture content of sediment samples was determined following ISO 11465 (gravimetric method: drying at 105°C for 24 h and mass difference). Although this standard originates from soil sciences, it has been applied successfully in marine contexts^18^, ensuring comparability and robustness of measurement.

**pH**

Sediment pH was measured using a protocol adapted from ISO 10390:2005. A 4 g portion of lyophilized sediment was placed into a 50 mL Falcon® tube, and 20 mL of deionized water were added. The suspension was mechanically shaken for 1 hour at 140 rpm and 20°C. Samples were then left to rest at room temperature for 1 to 3 hours before measurement. After manual homogenization and pH meter calibration, the pH value was recorded either after 2 minutes or once the reading had stabilized.

**Conductivity**

Sediment conductivity was assessed following a procedure inspired by ISO 11265:1994. A 20 g aliquot of lyophilized sediment was placed into a 200 mL plastic flask, to which 100 mL of deionized water were added. The suspension was then vortexed vigorously five times at 30-minute intervals over a 2.5-hour period. After agitation, the mixture was filtered using a Whatman No. 5 filter, and conductivity was measured on the resulting filtrate using a calibrated conductivity meter.

**Sediment grain size**

The sediment grain size distribution was measured using a Malvern laser diffraction instrument (LS13320XR), covering particle sizes from 0.4 to 2000 µm. Sediment samples (a few milligrams) were pretreated by removing organic matter (OM) with hydrogen peroxide and dispersed using sodium hexametaphosphate before analysis. The instrument’s high sensitivity allowed small sample volumes to be used, ensuring accurate and efficient granulometry. The distribution of particle sizes was expressed as the percentage of clay (<2 µm), silt (2-50 µm), and sand (50-2000 µm).

**Elementary analyser, nitrogen content**

Total nitrogen (N) content was quantified using elemental analysis (e.g. Carlo Erba NA-1500 CNS analyzer) on decarbonated, dried, unsalted and homogenized sediment samples. The analyzer combusts the material at high temperature (1200°C), allowing precise measurement of N as a percentage of dry weight.

**Nitrogen analyses**

For organic nitrogen pool analysis, fresh soil samples (5 g dry weight equivalent) were transferred into 150 mL plasma vials. After determining the soil moisture content, a 0.01 M CaCl₂ solution^19^ was added. The vials were sealed with parafilm (M®), and the soil suspensions were incubated at 20°C with shaking (140 rpm) for 2 hours at 10°C. Following incubation, the suspensions were filtered using membranes with a 0.22 μm pore size. Concentrations of NO₃⁻, NO₂⁻, and NH₄⁺ were determined using a Smartchem 200 sequential analyzer (AMS Alliance, Villeneuve-la-Garenne, France).

**Rock-Eval® Thermal analyses, hydrogen index**

Rock-Eval® thermal analysis is a geochemical technique widely used to quantify and characterize the carbon (C) content of sediment samples^20^. In this method, a small, dried, unsalted, and homogenized sediment sample undergoes two successive ramped heating steps: the first under an inert atmosphere (pyrolysis), followed by a second under an oxidative atmosphere.

During the pyrolysis phase, the sample is gradually heated from 200°C to 650°C. The thermal cracking of organic compounds releases a series of volatile products, mainly hydrocarbon effluents, carbon monoxide (CO), and carbon dioxide (CO₂), which are continuously monitored by flame ionization detector (FID) and infrared (IR) sensors. In the subsequent oxidation phase, the residual sample is heated from 200°C to 850°C in the presence of oxygen. This step enables the combustion of the remaining OM and the thermal decomposition of the mineral fraction, again releasing CO and CO₂, which are measured using IR detectors.

Based on the type, quantity, and temperature of gas release, Rock-Eval® allows the estimation of both total organic carbon (TOC [% g C·g⁻¹]) and mineral carbon (MinC [% g C·g⁻¹]). However, TOC values may sometimes be biased, particularly in non-carbonaceous samples, due to the misattribution of certain C signals, leading to an overestimation of the mineral C content. To correct for this analytical artifact, the SOTHIS correction methodwas applied to the entire dataset^21,22^. In the remainder of this article, *Cmin* refer to the MinC values corrected using the SOTHIS method.

In addition to quantitative C measurements, Rock-Eval® also provides qualitative indicators such as the Hydrogen Index (HI [mg HC·g⁻¹ TOC]). HI is calculated as the ratio between the quantity of hydrocarbons released during pyrolysis (S2 peak) and the TOC content (HI = S2/TOC × 100, in mg HC·g⁻¹ TOC). Higher HI values generally indicate hydrogen-rich, well-preserved (typically autochthonous) OM, while lower values reflect more degraded or terrestrially derived inputs.

**Analyses on the top 20 cm of undisturbed sediment cores**

Sediment cores (up to 20 cm depth) were manually collected using 35 cm long PVC tubes inserted until resistance. In the lab, the cores were sectioned at 1 cm intervals, dried at 62°C; wet and dry weights were used to calculate the dry bulk density (DBD), assuming a mineral density of 2.65 g cm⁻³. The Sediment Accumulation Rate (SAR) (cm yr⁻¹) and Mass Accumulation Rate (MAR) (g cm⁻² yr⁻¹) were calculated using excess ²¹⁰Pb (Pbₓₛ) profiles in the sediment, following Amann et al.,^23^. A Broad-Energy (@Mirion) gamma spectrometer was used to measure ²¹⁰Pb, ²²⁶Ra, and ²³²Th at the EPOC laboratory^24^. The excess of ^210^Pb (^210^Pb_ₓₛ_), incorporated in sediments from atmospheric inputs, whose decrease with depth provides access to time, was obtained by subtracting ²²⁶Ra from total ²¹⁰Pb and normalized using ²³²Th. Sediment ages were estimated using the Constant Initial Concentration model^25^. The SAR and MAR correspond to the slope of the depth-age and cumulative mass-age relationships, respectively.

C_org_ was measured in composite samples prepared from sediment layers located below the surface horizon affected by early diagenesis. Surface sediments were excluded to avoid bias due to OM degradation occurring within the uppermost centimeters. Following Delgard et al.,^26^, who showed in Arcachon Bay that beyond 3-5 cm depth %C remains relatively constant down to 20 cm, we considered this deeper section as the “stable interval.” Composite samples were prepared by pooling the selected layers within this interval for each core, with each layer weighted by its dry bulk density (DBD) to account for differences in sediment mass per unit volume. Sediments were ground, sieved (2 mm), and decarbonated following Lorrain et al.,^27^. C_org_ content was determined using a Thermo FlashSmart Elemental Analyzer at the EPOC laboratory. CAR (g C m⁻² yr⁻¹) was calculated as the product of MAR and %C_org_, and converted to CO₂ equivalents using a 44:12 molar ratio. The net climate benefit of each habitat was assessed by balancing CAR against GHG emissions. Further details on this method are available in Dolivet-Maréchal et al.,^3^. These values (SAR, MAR and CAR) were calculated for the sediment layer corresponding to the last 17-year, the recent period of sedimentary stabilization.

**Supplementary tables**

**Table S1. Target genes, primer names and sequences, amplicon sizes, and references used for quantitative PCR.**

| **Target Gene** | **Primer Name** | **Sequence (5’→3’)** | **Amplicon Size (bp)** | **Thermal Conditions** | **PCR Amplification Efficiency (%)** | **Standard Curve R²** | **Reference** |
| --- | --- | --- | --- | --- | --- | --- | --- |
| ***nirS*** | nirSCd3aF | GTSAACGTSAAGGARACSGG | 422 | Denaturation: 95°C/3min - 5 cycles: 95°C/15s; 59°C/30s - 35 cycles: 95°C/15s; 54°C/30s; 80°C/5s (data acquisition)  - Melting curve analysis: 0.5°C/0,05s from 65°C to 95°C | 83.8 | 0.998 | Throback et al. 2004 |
|  | nirSR3cd | GASTTCGGRTGSGTCTTGA |  |  |  |  |  |
| ***nirK*** | nirK876 | ATYGGCGGVCAYGGCGA | 164 | Denaturation: 95°C/3min - 5 cycles: 95°C/15s; 63°C/30s  - 30 cycles: 95°C/15s; 58°C/30s; 80°C/5s (data acquisition)  - Melting curve analysis: 0.5°C/0,05s from 65°C to 95°C | 90.7 | 0.999 | Henry et al. 2004 |
|  | nirK1040 | GCCTCGATCAGRTTRTGGTT |  |  |  |  |  |
| ***nosZ1*** | nosZ2F | CGCRACGGCAASAAGGTSMSSGT | 267 | Denaturation: 95°C/3min - 5 cycles: 95°C/15s; 65°C/30s  - 30 cycles: 95°C/15s; 60°C/30s; 80°C/5s (data acquisition)  - Melting curve analysis: 0.5°C/0,05s from 65°C to 95°C | 88.1 | 0.999 | Henry et al. 2006 |
|  | nosZ2R | CAKRTGCAKSGCRTGGCAGAA |  |  |  |  |  |
| ***nosZ2*** | nosZ-II-F | CTIGGICCIYTKCAYAC | 745 | Denaturation: 95°C/3min - 40 cycles: 95°C/15s; 58°C/60s; 80°C/5s (data acquisition)  - Melting curve analysis: 0.5°C/0,05s from 65°C to 95°C | 81 | 0.999 | Jones et al. 2013 |
|  | nosZ-II-R | GCIGARCARAAITCBGTRC |  |  |  |  |  |
| **16S rRNA** | 16S-341F | CCTACGGGNGGCWGCAG | 464 | Denaturation: 95°C/3min - 39 cycles: 95°C/15s; 60°C/1min; 80°C/5s (data acquisition) - Melting curve analysis: 0.5°C/0.05s from 65°C to 95°C | 81 | 0.997 | Klindworth et al. 2012 |
|  | 16S-805R | GACTACHVGGGTATCTAATCC |  |  |  |  |  |

**Table S2.** **Summary of significant mixed linear models explaining N_2_O fluxes across different variable subsets.** Models are ranked by Corrected Akaike Information Criterion (AICc), from lowest (best fit) to highest, and are grouped according to fixed or random factors: site, location, sediment type (Bare vs. Seagrass), subsite, Site nested within Type and Location nested within Type. All models use N_2_O flux as the response variable, with descriptors drawn from physical (A), chemical (B), microbial (C-I) variable categories. Only best models containing significant (p < 0.05) descriptors within each subset are presented. Variables included in each subset are listed at the top of their respective tables, n = 62.

A. **Physical**

| Subset of physical variables: Clay or Sand, Immersion_time, Hydro_forcing, Bathymetry, DBD, Moisture | | | | | |
| --- | --- | --- | --- | --- | --- |
| **Model** | **Formula: Fixed part** | **Random part** | **AICc (GLS)** | **Fixed R²** | **Global R²** |
| 1 | N_2_O_flux = - 0.46 x Clay - 0.39 x sqrt(Moisture) | ~1\|Type/Site | 148 | 0.41 | 0.71 |
| 2 | N_2_O_flux = - 0.45 x Clay - 0.42 x sqrt(Moisture) | ~1\|Type | 148 | 0.41 | 0.67 |
| 3 | N_2_O_flux = - 0.44 x Clay - 0.40 x sqrt(Moisture) | ~1\|Type/Location | 149 | 0.39 | 0.69 |
| 4 | N_2_O_flux = - 0.39 x Bathymetry - 0.35 x sqrt(Moisture) | ~1\|Subsite | 149 | 0.39 | 0.62 |
| 5 | N_2_O_flux = 0.57 x Sand | ~1\|Subsite | 150 | 0.31 | 0.60 |
| 6 | N_2_O_flux = - 0.56 x Immersion_time | ~1\|Subsite | 150 | 0.30 | 0.59 |
| 7 | N_2_O_flux = - 0.44 x sqrt(Moisture) | ~1\|Subsite | 150 | 0.21 | 0.57 |
| 8 | N_2_O_flux = - 0.55 x Clay | ~1\|Subsite | 151 | 0.28 | 0.60 |
| 9 | N_2_O_flux = - 0.54 x Bathymetry | ~1\|Subsite | 151 | 0.29 | 0.59 |
| 10 | N_2_O_flux = - 0.32 x sqrt(Moisture) | ~1\|Site | 155 | 0.12 | 0.44 |
| 11 | N_2_O_flux = - 0.40 x Hydro_forcing | ~1\|Site | 156 | 0.16 | 0.48 |
| 12 | N_2_O_flux = - 0.29 x sqrt(Moisture) | ~1\|Location | 157 | 0.08 | 0.46 |
| 13 | N_2_O_flux = 0.29 x Sand | ~1\|Location | 157 | 0.08 | 0.45 |
| 14 | N_2_O_flux = - 0.38 x Bathymetry - 0.38 x sqrt(Moisture) |  | 149(157) | 0.41 |  |
| 15 | N_2_O_flux = - 0.39 x Immersion_time + 0.35 x Sand |  | 149(157) | 0.41 |  |
| 16 | N_2_O_flux = 1.66 x Immersion_time - 1.28 x Bathymetry - 0.31 x sqrt(Moisture) | ~1\|Location | 158 | 0.06 | 0.79 |
| 17 | N_2_O_flux = 0.37 x Sand - 0.37 x Bathymetry |  | 150(158) | 0.40 |  |
| 18 | N_2_O_flux = - 0.39 x Immersion_time - 0.34 x sqrt(Moisture) |  | 150(158) | 0.40 |  |
| 19 | N_2_O_flux = - 0.28 x Clay - 0.28 x Bathymetry - 0.25 x Hydro_forcing |  | 150(160) | 0.42 |  |

B. **Chemical**

| Subset of chemical variables: pH, Conductivity, N, NH_4_, NO_2_, NO_3_, HI, C_min_, C_org_, CAR, SAR, MAR | | | | | |
| --- | --- | --- | --- | --- | --- |
| **Model** | **Formula: Fixed part** | **Random part** | **AICc (GLS)** | **Fixed R²** | **Global R²** |
| 1 | N_2_O_flux = 0.60 x MAR + 0.25 x pH | ~1\|Type/Site | 154 | 0.32 | 0.61 |
| 2 | N_2_O_flux = 0.52 x MAR | ~1\|Type/Site | 154 | 0.26 | 0.60 |
| 3 | N_2_O_flux = 0.44 x pH - 0.28 x HI | ~1\|Type/Location | 159 | 0.12 | 0.76 |
| 4 | N_2_O_flux = 0.31 x pH | ~1\|Type/Location | 159 | 0.07 | 0.63 |
| 5 | N_2_O_flux = 0.29 x pH - 0.19 x N | ~1\|Type/Location | 160 | 0.10 | 0.61 |
| 6 | N_2_O_flux = 0.58 x MAR + 0.27 x pH - 0.22 x N |  | 152(163) | 0.39 |  |
| 7 | N_2_O_flux = 0.41 x CAR - 0.41 x log_10_(Conductivity) - 0.27 x N + 0.21 x HI |  | 152(166) | 0.40 |  |
| 8 | N_2_O_flux = 0.58 x MAR + 0.27 x pH - 0.22 x N | ~1\|Type | 166 | 0.41 | 0.41 |
| 9 | N_2_O_flux = 0.41 x CAR - 0.29 x log_10_(Conductivity) - 0.27 x N + 0.21 x HI | ~1\|Type | 169 | 0.43 | 0.43 |

C. **Microbial**

| Subset of microbial variables: 16S, ratio_qPCR, DNA, Ratio_R, Ratio_D, *amoA*_AA_proxy, *amoA*_RA, *nrfA*_AA_proxy, *nrfA*_RA, *nirS*_D, *nirK*_D, *nosZ1*_D, *nosZ2*_D, *nirKS*, *nirKS*_AA, *nirKS*_RA, *nirKS*_R, *nosZ12*, *nosZ12*_AA, *nosZ12*_RA, *nosZ12*_R | | | | | |
| --- | --- | --- | --- | --- | --- |
| **Model** | **Formula: Fixed part** | **Random part** | **AICc (GLS)** | **Fixed R²** | **Global R²** |
| 1 | N_2_O_flux = 1.04 x *nosZ12* - 0.83 x *nirKS*_AA - 0.40 x *nirKS*_R | ~1\|Site | 150 | 0.12 | 0.71 |
| 2 | N_2_O_flux = 1.09 x *nosZ12* - 0.81 x *nirKS*_AA - 0.45 x *nirKS*_R - 0.22 x log_10_(DNA) | ~1\|Site | 150 | 0.17 | 0.70 |
| 3 | N_2_O_flux = 0.47 x *nosZ12* - 0.28 x *nirKS*_R | ~1\|Subsite | 153 | 0.14 | 0.63 |
| 4 | N_2_O_flux = 0.35 x *nrfA*_RA | ~1\|Location | 154 | 0.08 | 0.74 |
| 5 | N_2_O_flux = 0.49 x *nosZ12* - 0.35 x *nirKS*_R - 0.23 x log_10_(DNA) | ~1\|Subsite | 154 | 0.22 | 0.62 |
| 6 | N_2_O_flux = 0.29 x *nosZ12* | ~1\|Subsite | 154 | 0.09 | 0.59 |
| 7 | N_2_O_flux = 0.34 x *nrfA*_RA - 0.23 x log_10_(DNA) | ~1\|Location | 155 | 0.06 | 0.75 |
| 8 | N_2_O_flux = 0.45 x *nirKS* - 0.25 x *nirKS*_R | ~1\|Subsite | 155 | 0.14 | 0.61 |
| 9 | N_2_O_flux = - 0.93 x *nirKS*_AA + 0.80 x *nrfA*_AA_proxy - 0.26 x log_10_(DNA) | ~1\|Location | 156 | 0.08 | 0.74 |
| 10 | N_2_O_flux = 0.47 x *nosZ12* - 0.28 x *nirKS*_R | ~1\|Type/Site | 156 | 0.14 | 0.63 |
| 11 | N_2_O_flux = 0.29 x *nosZ12* | ~1\|Type/Site | 156 | 0.09 | 0.59 |
| 12 | N_2_O_flux = - 0.33 x log_10_(DNA) | ~1\|Type/Location | 156 | 0.10 | 0.56 |
| 13 | N_2_O_flux = 0.49 x *nosZ12* - 0.35 x *nirKS*_R - 0.23 x log_10_(DNA) | ~1\|Type/Site | 157 | 0.22 | 0.62 |
| 14 | N_2_O_flux = 0.45 x *nirKS* - 0.25 x *nirKS*_R | ~1\|Type/Site | 158 | 0.14 | 0.61 |
| 15 | N_2_O_flux = - 0.40 x log_10_(DNA) + 0.35 x *nirKS*_AA | ~1\|Type | 170 | 0.28 | 0.43 |
| 16 | N_2_O_flux = 0.51 x *nosZ12* - 0.43 x log_10_(DNA) - 0.29 x *nirKS*_R | ~1\|Type | 170 | 0.33 | 0.43 |
| 17 | N_2_O_flux = - 0.41 x log_10_(DNA) + 0.34 x *nosZ12* | ~1\|Type | 170 | 0.28 | 0.41 |
| 18 | N_2_O_flux = 0.49 x *nirKS*_AA - 0.41 x log_10_(DNA) - 0.25 x *nirKS*_R | ~1\|Type | 171 | 0.31 | 0.45 |
| 19 | N_2_O_flux = 0.48 x *nirKS* - 0.41 x log_10_(DNA) - 0.25 x *nirKS*_R | ~1\|Type | 171 | 0.31 | 0.41 |
| 20 | N_2_O_flux = - 0.40 x log_10_(DNA) + 0.33 x *nirKS* | ~1\|Type | 171 | 0.27 | 0.40 |
| 21 | N_2_O_flux = - 0.40 x log_10_(DNA) + 0.33 x log_10_(*nrfA*_AA_proxy) | ~1\|Type | 171 | 0.27 | 0.37 |
| 22 | N_2_O_flux = 0.55 x *nosZ12* - 0.36 x log_10_(DNA) - 0.32 x *nirKS*_R |  | 160(171) | 0.30 |  |
| 23 | N_2_O_flux = 0.52 x *nirKS* - 0.34 x log_10_(DNA) - 0.28 x *nirKS*_R |  | 162(172) | 0.29 |  |
| 24 | N_2_O_flux = 0.46 x *nirKS*_AA - 0.39 x log_10_(DNA) - 0.30 x *nirKS*_R - 0.24 x *nosZ12*_RA |  | 160(173) | 0.32 |  |

D. ***nirS* genus**

| Subset of *nirS* genus variables: tax9_NA, tax7_UBA5861, tax10_NA, tax2_*Thiogranum*, tax6_*Thiohalophilus*, tax16_*Sulfuriflexus*, tax25_GBMRC.2046, tax28_SZUA.150, tax26_*Prosthecomicrobium*_A, tax31_*Skermanella* | | | | | |
| --- | --- | --- | --- | --- | --- |
| **Model** | **Formula: Fixed part** | **Random part** | **AICc (GLS)** | **Fixed R²** | **Global R²** |
| 1 | N_2_O_flux = 0.90 x tax25_GBMRC.2046 - 0.74 x tax31_*Skermanella* | ~1\|Subsite | 155 | 0.08 | 0.57 |
| 2 | N_2_O_flux = 1.03 x tax25_GBMRC.2046 - 0.91 x tax31_*Skermanella* | ~1\|Site | 155 | 0.08 | 0.53 |
| 3 | N_2_O_flux = 1.03 x tax2_*Thiogranum* - 0.81 x tax10_NA | ~1\|Site | 156 | 0.12 | 0.51 |
| 4 | N_2_O_flux = 0.90 x tax25_GBMRC.2046 - 0.74 x tax31_*Skermanella* | ~1\|Type/Site | 155 | 0.08 | 0.57 |
| 5 | N_2_O_flux = 1.67 x tax6_*Thiohalophilus* - 1.22 x tax10_NA | ~1\|Type | 167 | 0.29 | 0.36 |
| 6 | N_2_O_flux = 0.17 x tax6_*Thiohalophilus* - 0.12 x tax10_NA |  | 162(167) | 0.27 |  |

E. ***nirK* genus**

| Subset of *nirK* genus variables: tax5_WVYX01, tax10_*Nitrosotenuis*, tax12_*Marivita*, tax22_*Sulfitobacter*, tax26_*Marivita*, tax15_*Crocosphaera*, tax25_*Planktotalea*, tax13_*Shimia*, tax9_*Nitrosopumilus*, tax3_*Nitrososphaera* | | | | | |
| --- | --- | --- | --- | --- | --- |
| **Model** | **Formula: Fixed part** | **Random part** | **AICc (GLS)** | **Fixed R²** | **Global R²** |
| 1 | N_2_O_flux = 0.31 x tax22_*Sulfitobacter* | ~1\|Subsite | 153 | 0.10 | 0.56 |
| 2 | N_2_O_flux = 0.25 x tax25_*Planktotalea* | ~1\|Subsite | 154 | 0.07 | 0.56 |
| 3 | N_2_O_flux = 0.33 x tax22_*Sulfitobacter* | ~1\|Site | 154 | 0.11 | 0.49 |
| 4 | N_2_O_flux = 0.31 x tax22_*Sulfitobacter* | ~1\|Type/Site | 155 | 0.10 | 0.56 |
| 5 | N_2_O_flux = 0.25 x tax25_*Planktotalea* | ~1\|Type/Site | 156 | 0.07 | 0.56 |
| 6 | N_2_O_flux = 0.26 x tax25_*Planktotalea* | ~1\|Site | 156 | 0.07 | 0.49 |
| 7 | N_2_O_flux = 0.49 x tax22_*Sulfitobacter* | ~1\|Type | 171 | 0.23 | 0.27 |
| 8 | N_2_O_flux = 0.48 x tax12_*Marivita* | ~1\|Type | 172 | 0.22 | 0.27 |
| 9 | N_2_O_flux = 0.49 x tax22_*Sulfitobacter* |  | 164(169) | 0.23 |  |
| $10 | N_2_O_flux = 0.48 x tax12_*Marivita* |  | 165(170) | 0.22 |  |

F. ***nosZ1* genus**

| Subset of *nosZ1* genus variables: tax1_NA, tax5_*Rhodobacteraceae_bacterium*, tax9_*Emcibacter*, tax10_*Roseobacter*, tax25_NA, tax14_NA, tax15_*Thalassospira*,tax21_*Ruegeria*, tax20_NA, tax36_*Pseudophaeobacter* | | | | | |
| --- | --- | --- | --- | --- | --- |
| **Model** | **Formula: Fixed part** | **Random part** | **AICc (GLS)** | **Fixed R²** | **Global R²** |
| 1 | N_2_O_flux = 0.72 x tax1_NA - 0.47 x tax10_*Roseobacter* | ~1\|Subsite | 152 | 0.20 | 0.67 |
| 2 | N_2_O_flux = 0.27 x tax25_NA | ~1\|Subsite | 152 | 0.08 | 0.57 |
| 3 | N_2_O_flux = 0.98 x tax1_NA - 0.49 x tax10_*Roseobacter* - 0.26 x tax20_NA | ~1\|Subsite | 153 | 0.25 | 0.71 |
| 4 | N_2_O_flux = 0.73 x tax1_NA - 0.48 x tax10_*Roseobacter* | ~1\|Type/Site | 154 | 0.20 | 0.68 |
| 5 | N_2_O_flux = 0.27 x tax9_*Emcibacter* | ~1\|Subsite | 154 | 0.08 | 0.55 |
| 6 | N_2_O_flux = 0.27 x tax25_NA | ~1\|Type/Site | 155 | 0.08 | 0.57 |
| 7 | N_2_O_flux = 0.28 x tax9_*Emcibacter* | ~1\|Site | 155 | 0.08 | 0.48 |
| 8 | N_2_O_flux = 0.27 x tax9_*Emcibacter* | ~1\|Type/Site | 156 | 0.08 | 0.55 |
| 9 | N_2_O_flux = 0.32 x tax1_NA |  | 164(169) | 0.09 |  |
| 10 | N_2_O_flux = 0.31 x tax9_*Emcibacter* + 0.29 x tax25_NA |  | 162(170) | 0.26 |  |
| 11 | N_2_O_flux = 0.49 x tax25_NA | ~1\|Type | 171 | 0.23 | 0.30 |
| 12 | N_2_O_flux = 0.49 x tax1_NA | ~1\|Type | 171 | 0.23 | 0.26 |
| 13 | N_2_O_flux = 0.47 x tax5_*Rhodobacteraceae_bacterium* | ~1\|Type | 172 | 0.22 | 0.26 |
| 14 | N_2_O_flux = 0.47 x tax5_*Rhodobacteraceae_bacterium* | ~1\|Type | 172 | 0.22 | 0.26 |
| 15 | N_2_O_flux = 0.49 x tax9_*Emcibacter* | ~1\|Type | 172 | 0.23 | 0.23 |
| 16 | N_2_O_flux = 0.33 x tax9_*Emcibacter* + 0.28 x tax14_NA | ~1\|Type | 173 | 0.26 | 0.31 |

G. ***nosZ2* genus**

| Subset of *nosZ2* genus variables: tax9_NA, tax10_NA, tax13_NA, tax16_NA, tax7_NA, tax20_*Robiginitalea*, tax26_NA, tax30_*Flavobacteriaceae_bacterium*, tax18_*Muricauda*, tax23_*Gramella* | | | | | |
| --- | --- | --- | --- | --- | --- |
| **Model** | **Formula: Fixed part** | **Random part** | **AICc gGLS)** | **Fixed R²** | **Global R²** |
| 1 | N_2_O_flux = - 0.61 x tax10_NA + 0.54 x tax16_NA | ~1\|Type/Location | 163 | 0.05 | 0.59 |
| 2 | N_2_O_flux = 0.76 x tax26_NA - 0.75 x tax7_NA + 0.34 x tax18_*Muricauda* |  | 165(173) | 0.25 |  |
| 3 | N_2_O_flux = - 0.77 x tax7_NA + 0.70 x tax26_NA + 0.45 x tax13_NA | ~1\|Type | 173 | 0.26 | 0.43 |
| 4 | N_2_O_flux = 0.75 x tax26_NA - 0.74 x tax7_NA + 0.35 x tax18_*Muricauda* | ~1\|Type | 174 | 0.27 | 0.33 |
| 5 | N_2_O_flux = - 0.77 x tax7_NA + 0.71 x tax26_NA + 0.40 x tax20_*Robiginitalea* | ~1\|Type | 175 | 0.26 | 0.35 |
| 6 | N_2_O_flux = 0.99 x tax26_NA - 0.73 x tax7_NA | ~1\|Type | 175 | 0.22 | 0.27 |

H. **Network variables by type**

| Subset of network_type variables: mean_efficiency_type, Vertex_type, Edge_type, Average_degree_type, Average_path_length_type, Clustering_coefficient_type, Density_type, Heterogeneity_type, Centralization_type, Modularity_type, nat_co_type, Percent_Pos_Edges_type, Percent_Neg_Edges_type, mean_vulnerability_type, assortativity_type | | | | | |
| --- | --- | --- | --- | --- | --- |
| **Model** | **Formula: Fixed part** | **Random part** | **AICc (GLS)** | **Fixed R²** | **Global R²** |
| 1 | N_2_O_flux = - 1.72 x Edge_type + 1.60 x Average_degree_type | ~1\|Site | 155 | 0.05 | 0.58 |
| 2 | N_2_O_flux = 0.29 x Density_type |  | 176(181) | 0.07 |  |
| 3 | N_2_O_flux = 1.08 x Average_degree_type - 1.02 x Vertex_type |  | 178(182) | 0.06 |  |
| 4 | N_2_O_flux = 0.29 x Density_type | ~1\|Type | 183 | 0.08 | 0.08 |
| 5 | N_2_O_flux = - 2.02 x Edge_type + 2.10 x Average_degree_type | ~1\|Site | 184 | 0.07 | 0.07 |

I. **Network variables by location**

| Subset of network_location variables: mean_efficiency_location, Vertex_location, Edge_location, Average_degree_location, Average_path_length_location, Clustering_coefficient_location, Density_location, Heterogeneity_location, Centralization_location, Modularity_location, nat_co_location, Percent_Pos_Edges_location, Percent_Neg_Edges_location, mean_vulnerability_location, assortativity_location | | | | | |
| --- | --- | --- | --- | --- | --- |
| **Model** | **Formula: Fixed part** | **Random part** | **AICc (GLS)** | **Fixed R²** | **Global R²** |
| 1 | N_2_O_flux = - 0.56 x Heterogeneity_location | ~1\|Subsite | 147 | 0.32 | 0.60 |
| 2 | N_2_O_flux = 0.58 x Clustering_coefficient_location - 0.39 x Average_path_length_location | ~1\|Subsite | 147 | 0.45 | 0.66 |
| 3 | N_2_O_flux = - 0.56 x Heterogeneity_location | ~1\|Type/Site | 149 | 0.32 | 0.61 |
| 4 | N_2_O_flux = 0.58 x Clustering_coefficient_location - 0.39 x Average_path_length_location | ~1\|Type/Site | 149 | 0.45 | 0.66 |
| 5 | N_2_O_flux = - 0.44 x mean_efficiency_location + 0.38 x Clustering_coefficient_location + 0.36 x mean_vulnerability_location | ~1\|Site | 151 | 0.49 | 0.56 |
| 6 | N_2_O_flux = 0.63 x Clustering_coefficient_location - 0.36 x Average_path_length_location | ~1\|Site | 151 | 0.46 | 0.60 |
| 7 | N_2_O_flux = - 0.41 x Heterogeneity_location - 0.25 x mean_efficiency_location | ~1\|Site | 151 | 0.36 | 0.52 |
| 8 | N_2_O_flux = - 0.53 x Heterogeneity_location | ~1\|Site | 151 | 0.29 | 0.50 |
| 9 | N_2_O_flux = - 0.51 x mean_efficiency_location + 0.39 x mean_vulnerability_location + 0.34 x Clustering_coefficient_location |  | 139(151) | 0.50 |  |
| 10 | N_2_O_flux = - 0.57 x mean_efficiency_location + 0.48 x mean_vulnerability_location - 0.30 x assortativity_location |  | 140(152) | 0.50 |  |
| 11 | N_2_O_flux = 0.46 x Clustering_coefficient_location - 0.35 x Average_path_length_location - 0.25 x mean_efficiency_location | ~1\|Site | 152 | 0.50 | 0.61 |
| 12 | N_2_O_flux = - 0.42 x mean_efficiency_location - 0.37 x Heterogeneity_location + 0.24 x mean_vulnerability_location | ~1\|Site | 152 | 0.46 | 0.56 |
| 13 | N_2_O_flux = - 0.49 x mean_efficiency_location + 0.39 x mean_vulnerability_location + 0.34 x Clustering_coefficient_location | ~1\|Type | 153 | 0.51 | 0.53 |
| 14 | N_2_O_flux = - 0.54 x mean_efficiency_location + 0.45 x mean_vulnerability_location - 0.33 x assortativity_location | ~1\|Type | 153 | 0.50 | 0.53 |
| 15 | N_2_O_flux = - 0.32 x mean_efficiency_location | ~1\|Site | 153 | 0.11 | 0.47 |
| 16 | N_2_O_flux = - 0.50 x mean_efficiency_location + 0.40 x mean_vulnerability_location - 0.30 x assortativity_location | ~1\|Site | 153 | 0.46 | 0.52 |
| 17 | N_2_O_flux = 1.69 x Modularity_location + 1.27 x Average_degree_location - 0.59 x Average_path_length_location - 0.37 x Heterogeneity_location - 0.29 x mean_efficiency_location | ~1\|Site | 153 | 0.47 | 0.59 |
| 18 | N_2_O_flux = 1.64 x Modularity_location + 1.09 x Edge_location - 0.68 x Average_path_length_location - 0.46 x Heterogeneity_location - 0.29 x mean_efficiency_location | ~1\|Site | 153 | 0.47 | 0.59 |
| 19 | N_2_O_flux = - 0.51 x mean_efficiency_location + 0.39 x mean_vulnerability_location + 0.34 x Clustering_coefficient_location | ~1\|Location | 153 | 0.52 | 0.52 |
| 20 | N_2_O_flux = - 0.42 x Average_path_length_location - 0.38 x assortativity_location - 0.30 x mean_efficiency_location | ~1\|Type | 154 | 0.47 | 0.55 |
| 21 | N_2_O_flux = - 0.48 x mean_efficiency_location - 0.33 x Heterogeneity_location + 0.29 x mean_vulnerability_location | ~1\|Type | 154 | 0.49 | 0.53 |
| 22 | N_2_O_flux = 0.38 x Clustering_coefficient_location - 0.35 x Average_path_length_location - 0.29 x mean_efficiency_location | ~1\|Type | 154 | 0.48 | 0.53 |
| 23 | N_2_O_flux = - 0.34 x mean_efficiency_location | ~1\|Location | 154 | 0.11 | 0.49 |
| 24 | N_2_O_flux = - 0.55 x mean_efficiency_location + 0.28 x mean_vulnerability_location | ~1\|Location | 154 | 0.28 | 0.44 |
| 25 | N_2_O_flux = - 0.57 x mean_efficiency_location + 0.45 x mean_vulnerability_location - 0.30 x assortativity_location | ~1\|Location | 154 | 0.51 | 0.51 |
| 26 | N_2_O_flux = 0.82 x Modularity_location - 0.73 x Average_path_length_location - 0.38 x mean_efficiency_location | ~1\|Location | 155 | 0.09 | 0.67 |
| 27 | N_2_O_flux = - 0.52 x mean_efficiency_location + 0.31 x mean_vulnerability_location - 0.29 x mean_efficiency_location | ~1\|Location | 155 | 0.50 | 0.50 |
| 28 | N_2_O_flux = - 0.49 x mean_efficiency_location + 0.39 x mean_vulnerability_location + 0.35 x Clustering_coefficient_location | ~1\|Type/Location | 155 | 0.51 | 0.53 |
| 29 | N_2_O_flux = - 0.54 x mean_efficiency_location + 0.45 x mean_vulnerability_location - 0.33 x assortativity_location | ~1\|Type/Location | 156 | 0.50 | 0.53 |
| 30 | N_2_O_flux = - 0.42 x Average_path_length_location - 0.38 x assortativity_location - 0.30 x mean_efficiency_location | ~1\|Type/Location | 156 | 0.48 | 0.55 |
| 31 | N_2_O_flux = - 0.48 x mean_efficiency_location - 0.33 x Heterogeneity_location + 0.29 x mean_vulnerability_location | ~1\|Type/Location | 157 | 0.49 | 0.53 |
| 32 | N_2_O_flux = 0.38 x Clustering_coefficient_location - 0.35 x Average_path_length_location - 0.29 x mean_efficiency_location | ~1\|Type/Location | 157 | 0.48 | 0.53 |

| **Site** | **Site_ID** | **Type** | **Location** | **Longitude** | **Latitude** | **Plant_cover** |  | **Sediment texture** | **Clay** |  | **Silt** |  | **Sand** |  |
| --- | --- | --- | --- | --- | --- | --- | --- | --- | --- | --- | --- | --- | --- | --- |
|  |  | **(Subsite)** |  | **°E. WEGS84** | **°N. WGS84** |  |  |  | **%** |  | **%** |  | **%** |  |
| Germanan | Germ | Bare | Outer | -1.13384771347 | 44.7116661072 | 0 |  | Sandy silt | 4.1 |  | 63.4 |  | 32.5 |  |
|  |  | Seagrass |  | -1.13384771347 | 44.7116661072 | 0.9 |  | Sandy silt | 4.1 |  | 55.4 |  | 40.5 |  |
| Comprian | Comp | Bare | Outer | -1.070611 | 44.679473 | 0 |  | Silty sand | 3.0 |  | 47.4 |  | 49.6 |  |
|  |  | Seagrass |  | -1.072061 | 44.679063 | 0.9 |  | Silty sand | 3.8 |  | 47.8 |  | 48.4 |  |
| Garrèche | Garr | Bare | Outer | -1.1205807 | 44.7047449 | 0 |  | Silty sand | 1.7 |  | 26 |  | 72.3 |  |
|  |  | Seagrass |  | -1.1208349 | 44.7040884 | 0.4 |  | Silty sand | 2.5 |  | 37.8 |  | 59.7 |  |
| Gaillard | Gail | Bare | Outer | -1.0994219 | 44.6627752 | 0 |  | Sandy silt | 3.5 |  | 61.2 |  | 35.3 |  |
|  |  | Seagrass |  | -1.09981 | 44.66241 | 0.9 |  | Sandy silt | 3.2 |  | 60.4 |  | 36.4 |  |
| Andernos | Ande | Bare | Inner | -1.12038 | 44.74512 | 0 |  | Sand | 1.3 |  | 23.7 |  | 75.0 |  |
|  |  | Seagrass |  | -1.12138 | 44.74513 | 0.9 |  | Sand | 1.3 |  | 21.5 |  | 77.2 |  |
| Fontaines | Font | Bare | Inner | -1.07578 | 44.71885 | 0 |  | Sand | 0.6 |  | 8.4 |  | 91.0 |  |
|  |  | Seagrass |  | -1.0754 | 44.71874 | 0.9 |  | Sandy silt | 3.6 |  | 52.4 |  | 44.0 |  |
| All sites | All | Bare |  | NA | NA | 0.0 | *** | NA | 2.4 ± 0.2 | * | 38.1 ± 3.5 | NS | 59.6 ± 3.8 | NS |
|  |  | Seagrass |  |  |  | 0.8 ± 0.0 |  |  | 3.1 ± 0.2 |  | 45.9 ± 2.2 |  | 51.1 ± 2.3 |  |
| Location | Inner |  |  | NA | NA | 0.42 ± 0.09 | NS | NA | 1.7 ± 0.2 | *** | 26.5 ± 3.4 | *** | 71.8 ± 3.6 | *** |
|  | Outer |  |  |  |  | 0.38 ± 0.06 |  |  | 3.2 ± 0.1 |  | 50.0 ± 1.8 |  | 46.8 ± 1.9 |  |

**Table S3. Characteristics of the studied sites under Bare and vegetated (Seagrass) conditions, with variables grouped into general and physical features (A) and chemical variables (B).** Significant differences (Wilcoxon tests) between Bare and Seagrass sediments, and Inner and Outer sites are indicated as follows: # (p < 0.1), * (p < 0.05), ** (p < 0.01) ), *** (p < 0.001), and NS (not significant), n = 71.

**A**

| **Site** | **Site_ID** | **Type** | **Location** |  | **Bathymetry** |  | **Hydrodynamic forcing** |  | **Immersion_time** |  | **DBD** |  | **Moisture** |  |
| --- | --- | --- | --- | --- | --- | --- | --- | --- | --- | --- | --- | --- | --- | --- |
|  |  | **(Subsite)** |  |  | **m** |  |  |  | **h day⁻¹** |  | **g cm⁻³** |  | **%** |  |
| Germanan | Germ | Bare | Outer |  | 1.7 |  | 1.3e+05 |  | 18.0 |  | 0.8 |  | 62.8 ± 3.5 | NS |
|  |  | Seagrass |  |  | 1.7 |  | 1.3e+05 |  | 18.0 |  | 1.2 |  | 58.1 ± 4.5 |  |
| Comprian | Comp | Bare | Outer |  | 1.4 |  | 2.1e+05 |  | 18.3 |  | 1.1 |  | 39.3 ± 3.0 | ** |
|  |  | Seagrass |  |  | -0.2 |  | 8.3e+04 |  | 10.6 |  | 1.0 |  | 55.0 ± 3.4 |  |
| Garrèche | Garr | Bare | Outer |  | 1.2 |  | 2.3e+05 |  | 17.3 |  | 1.1 |  | 52.7 ± 5.3 | NS |
|  |  | Seagrass |  |  | 1.1 |  | 1.1e+05 |  | 17.3 |  | 1.1 |  | 56.7 ± 7.3 |  |
| Gaillard | Gail | Bare | Outer |  | 0.3 |  | 3.9e+05 |  | 13.1 |  | 0.8 |  | 79.8 ± 3.6 | NS |
|  |  | Seagrass |  |  | 0.5 |  | 2.8e+05 |  | 13.1 |  | 0.8 |  | 74.0 ± 5.5 |  |
| Andernos | Ande | Bare | Inner |  | -0.8 |  | 0 |  | 1.7 |  | 1.4 |  | 34.8 ± 1.3 | ** |
|  |  | Seagrass |  |  | -0.5 |  | 0 |  | 5.6 |  | 1.1 |  | 52.1 ± 2.7 |  |
| Fontaines | Font | Bare | Inner |  | -1.1 |  | 0 |  | 1.7 |  | 1.7 |  | 23.2 ± 0.5 | ** |
|  |  | Seagrass |  |  | -1.1 |  | 0 |  | 1.7 |  | 1.2 |  | 38.5 ± 2.3 |  |
| All sites | All | Bare |  | NS | 0.4 ± 0.2 | NS | 1.6e+05 ± 2.4e+04 | # | 11.5 ± 1.3 | NS | 1.2 ± 0.1 | NS | 49.1 ± 3.5 | NS |
|  |  | Seagrass |  |  | 0.3 ± 0.2 |  | 9.9e+04 ± 1.6e+04 |  | 11.1 ± 1.0 |  | 1.1 ± 0.0 |  | 55.7 ± 2.5 |  |
| Location | Inner |  |  | *** | -0.9 ± 0.1 | *** | 0 ± 0 | *** | 2.7 ± 0.4 | *** | 1.3 ± 0.1 | *** | 37.1 ± 2.3 | *** |
|  | Outer |  |  |  | 1.0 ± 0.1 |  | 1.9e+05 ± 1.4e+04 |  | 15.7 ± 0.4 |  | 1.0 ± 0.0 |  | 60.2 ± 2.3 |  |

**B**

| **Site_ID** | **Type** | **Location** | **pH** |  | **Conductivity** |  | **C_mineral_** |  | **C_organic_** |  | **HI** |  |
| --- | --- | --- | --- | --- | --- | --- | --- | --- | --- | --- | --- | --- |
|  | **(Subsite)** |  |  |  | **mS cm⁻¹** |  | **%** |  | **%** |  |  |  |
| Germ | Bare | Outer | 7.3 ± 0.6 | NS | 6.6 ± 0.5 | NS | 0.158 ± 0.089 | NS | 1.5 |  | 208.2 ± 17.4 | ** |
|  | Seagrass |  | 8.1 ± 0.2 |  | 6.0 ± 0.8 |  | 0.067 ± 0.004 |  | 1.5 |  | 246.7 ± 3.6 |  |
| Comp | Bare | Outer | 8.5 ± 0.0 | NS | 3.5 ± 0.3 | NS | 0.082 ± 0.018 | * | 1.1 |  | 190.6 ± 3.0 | NS |
|  | Seagrass |  | 8.5 ± 0.0 |  | 3.2 ± 0.2 |  | 0.360 ± 0.095 |  | 0.6 |  | 201.0 ± 5.2 |  |
| Garr | Bare | Outer | 8.4 ± 0.1 | NS | 5.9 ± 0.6 | NS | 0.751 ± 0.090 | NS | 1.3 |  | 188.3 ± 7.8 | NS |
|  | Seagrass |  | 8.4 ± 0.1 |  | 6.4 ± 0.6 |  | 0.646 ± 0.184 |  | 0.6 |  | 201.8 ± 3.6 |  |
| Gail | Bare | Outer | 8.0 ± 0.0 | NS | 7.4 ± 1.0 | NS | 0.164 ± 0.015 | * | 0.9 |  | 229.0 ± 2.2 | NS |
|  | Seagrass |  | 7.4 ± 0.4 |  | 6.6 ± 1.0 |  | 0.107 ± 0.019 |  | 2.0 |  | 209.3 ± 13.1 |  |
| Ande | Bare | Inner | 6.8 ± 0.5 | * | 4.0 ± 0.2 | ** | 0.011 ± 0.003 | ** | 0.5 |  | 250.7 ± 12.3 | # |
|  | Seagrass |  | 7.8 ± 0.1 |  | 6.2 ± 0.4 |  | 0.081 ± 0.029 |  | 1.1 |  | 242.8 ± 4.4 |  |
| Font | Bare | Inner | 6.8 ± 0.1 | ** | 4.4 ± 1.2 | # | 0.020 ± 0.004 | * | 0.1 |  | 243.0 ± 10.6 | NS |
|  | Seagrass |  | 8.4 ± 0.1 |  | 4.3 ± 0.3 |  | 0.096 ± 0.022 |  | 0.8 |  | 240.5 ± 1.4 |  |
| All | Bare |  | 7.6 ± 0.2 | * | 5.3 ± 0.4 | NS | 0.201 ± 0.049 | NS | 0.9 ± 0.1 | NS | 219.1 ± 5.8 | NS |
|  | Seagrass |  | 8.1 ± 0.1 |  | 5.5 ± 0.3 |  | 0.226 ± 0.049 |  | 1.1 ± 0.1 |  | 223.7 ± 4.1 |  |
| Location | Inner |  | 7.5 ± 0.2 | *** | 4.7 ± 0.4 | * | 0.052 ± 0.012 | *** | 0.6 ± 0.1 | *** | 244.3 ± 4.0 | *** |
|  | Outer |  | 8.1 ± 0.1 |  | 5.8 ± 0.3 |  | 0.322 ± 0.040 |  | 1.2 ± 0.1 |  | 209.8 ± 4.0 |  |

| **Site_ID** | **Type** | **Location** | **SAR** |  | **MAR** |  | **CAR** |  | **Climate regulation** |  |
| --- | --- | --- | --- | --- | --- | --- | --- | --- | --- | --- |
|  | **(Subsite)** |  | **cm y⁻¹** |  | **g cm⁻² y⁻¹** |  | **g CO₂-eq m⁻² y⁻¹** |  | **g CO₂-eq m⁻² y⁻¹** |  |
| Germ | Bare | Outer | 0.28 |  | 0.13 |  | 38.8 |  | 48.5 ± 1.7 | ** |
|  | Seagrass |  | 0.11 |  | 0.09 |  | 22.9 |  | 18.6 ± 1.6 |  |
| Comp | Bare | Outer | 0.09 |  | 0.10 |  | 33.8 |  | 21.6 ± 2.8 | ** |
|  | Seagrass |  | 0.09 |  | 0.06 |  | 12.6 |  | 5.1 ± 1.1 |  |
| Garr | Bare | Outer | 0 |  | 0 |  | 0 |  | -13.0 ± 5.1 | ** |
|  | Seagrass |  | 0.32 |  | 0.33 |  | 92.2 |  | 78.9 ± 3.0 |  |
| Gail | Bare | Outer | 0 |  | 0 |  | 0 |  | 5.2 ± 1.8 | ** |
|  | Seagrass |  | 0.15 |  | 0.11 |  | 62.6 |  | 47.8 ± 1.7 |  |
| Ande | Bare | Inner | 0.29 |  | 0.40 |  | 74.5 |  | 55.8 ± 5.3 | ** |
|  | Seagrass |  | 0.35 |  | 0.28 |  | 129.9 |  | 104.0 ± 1.6 |  |
| Font | Bare | Inner | 0.23 |  | 0.37 |  | 21.7 |  | 2.3 ± 2.2 | ** |
|  | Seagrass |  | 0.26 |  | 0.30 |  | 78.9 |  | 53.9 ± 4.4 |  |
| All | Bare |  | 0.15 ± 0.02 | * | 0.17 ± 0.03 | NS | 27.9 ± 4.4 | *** | 20.0 ± 4.5 | *** |
|  | Seagrass |  | 0.21 ± 0.02 |  | 0.20 ± 0.02 |  | 66.6 ± 6.8 |  | 51.4 ± 5.8 |  |
| Location | Inner |  | 0.28 ± 0.1 | *** | 0.34 ± 0.01 | *** | 76.2 ± 7.9 | *** | 54.0 ± 7.7 | *** |
|  | Outer |  | 0.13 ± 0.02 |  | 0.10 ± 0.01 |  | 32.9 ± 4.4 |  | 26.7 ± 4.3 |  |

| **Site_ID** | **Type** | **Location** | **N** |  | **NH₄⁺** |  | **NO₂⁻** |  | **NO₃⁻** |  | **ratio_CN** |  |
| --- | --- | --- | --- | --- | --- | --- | --- | --- | --- | --- | --- | --- |
|  | **(Subsite)** |  | **%** |  | **µg N-NH₄ g⁻¹ dry soil** |  | **µg N-NO_2_ g⁻¹ dry soil** |  | **µg N-NO_3_ g⁻¹ dry soil** |  |  |  |
| Germ | Bare | Outer | 0.139 ± 0.009 | NS | 6.6 ± 1.0 | # | 0.014 ± 0.003 | NS | 0.105 ± 0.101 | # | 11.399 ± 0.910 | NS |
|  | Seagrass |  | 0.129 ± 0.014 |  | 8.1 ± 0.5 |  | 0.015 ± 0.002 |  | 0.439 ± 0.122 |  | 12.056 ± 1.335 |  |
| Comp | Bare | Outer | 0.070 ± 0.008 | ** | 19.7 ± 5.3 | NS | 0.020 ± 0.003 | NS | 0.450 ± 0.213 | NS | 16.274 ± 2.344 | ** |
|  | Seagrass |  | 0.119 ± 0.011 |  | 10.6 ± 2.1 |  | 0.019 ± 0.001 |  | 0.587 ± 0.088 |  | 5.037 ± 0.385 |  |
| Garr | Bare | Outer | 0.116 ± 0.009 | NS | 5.9 ± 0.4 | NS | 0.020 ± 0.002 | NS | 0.253 ± 0.160 | NS | 11.589 ± 0.846 | ** |
|  | Seagrass |  | 0.118 ± 0.009 |  | 6.3 ± 0.7 |  | 0.021 ± 0.010 |  | 0.431 ± 0.208 |  | 5.456 ± 0.390 |  |
| Gail | Bare | Outer | 0.202 ± 0.009 | # | 2.5 ± 0.4 | # | 0.016 ± 0.001 | NS | 0.072 ± 0.014 | NS | 4.296 ± 0.182 | ** |
|  | Seagrass |  | 0.158 ± 0.017 |  | 5.0 ± 0.8 |  | 0.018 ± 0.001 |  | 0.152 ± 0.081 |  | 13.498 ± 1.739 |  |
| Ande | Bare | Inner | 0.155 ± 0.084 | # | 2.8 ± 0.8 | * | 0.020 ± 0.009 | NS | 0.064 ± 0.016 | NS | 5.647 ± 0.988 | ** |
|  | Seagrass |  | 0.107 ± 0.015 |  | 5.4 ± 1.1 |  | 0.017 ± 0.001 |  | 0.082 ± 0.010 |  | 10.831 ± 1.324 |  |
| Font | Bare | Inner | 0.024 ± 0.008 | ** | 4.3 ± 0.8 | # | 0.009 ± 0.001 | ** | 0.219 ± 0.031 | NS | 9.074 ± 2.160 | NS |
|  | Seagrass |  | 0.067 ± 0.011 |  | 6.8 ± 1.3 |  | 0.017 ± 0.002 |  | 0.163 ± 0.017 |  | 13.860 ± 3.278 |  |
| All | Bare |  | 0.119 ± 0.017 | NS | 6.6 ± 1.2 | * | 0.016 ± 0.002 | NS | 0.186 ± 0.047 | * | 9.526 ± 0.848 | NS |
|  | Seagrass |  | 0.116 ± 0.007 |  | 7.1 ± 0.6 |  | 0.018 ± 0.002 |  | 0.309 ± 0.052 |  | 10.123 ± 0.887 |  |
| Location | Inner |  | 0.088 ± 0.023 | *** | 4.8 ± 0.6 | * | 0.016 ± 0.002 | * | 0.132 ± 0.016 | NS | 9.853 ± 1.170 | NS |
|  | Outer |  | 0.130 ± 0.006 |  | 7.9 ± 0.9 |  | 0.017 ± 0.001 |  | 0.311 ± 0.045 |  | 9.820 ± 0.710 |  |

**Table S4. Percentage of unknown taxa for the four studied genes at the genus and species taxonomic levels.**
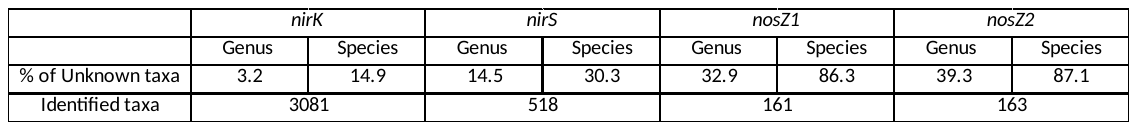

**Supplemetary figures**

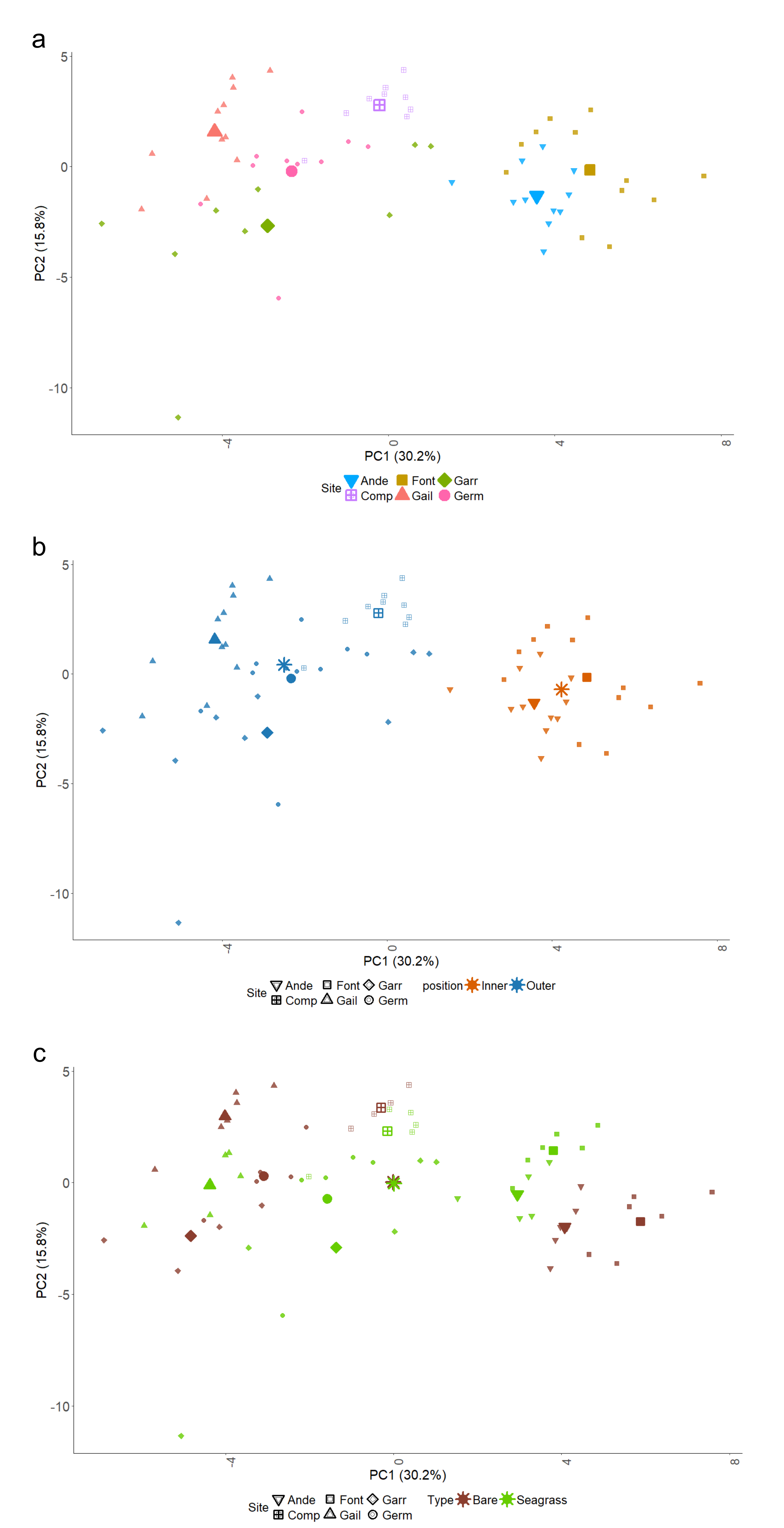

**Figure S1.** PCA of physical, chemical, microbial, and GHG variables across all sampling sites. Only sample scores are shown to illustrate the relative spatial arrangement of samples along PC1 and PC2. (**a)** Colors and point shapes indicate sampling sites. (**b)** Colors indicate bay location (Inner vs. Outer), while point shapes indicate sampling sites. (**c)** Colors indicate vegetation status (Seagrass vs. Bare), while point shapes indicate sampling sites.

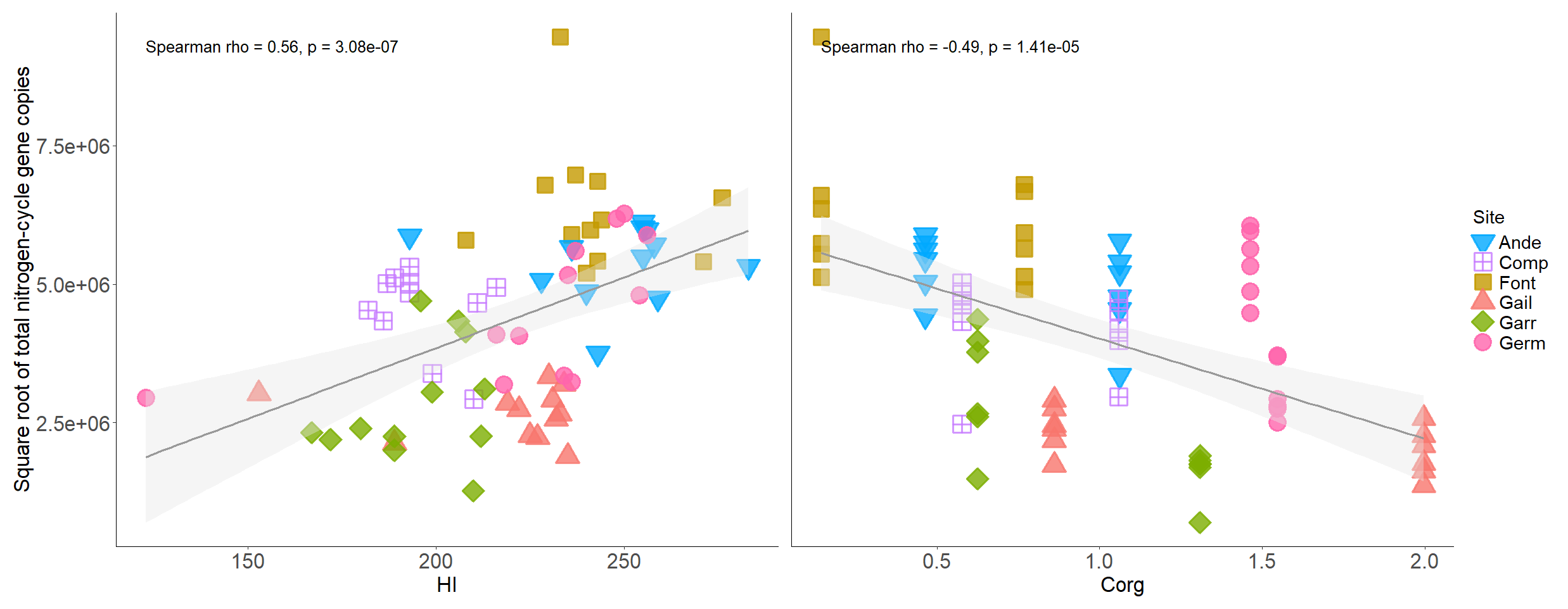

**Figure S2.** Relationships between microbial abundance and sediment characteristics across all sites. Left panel: square root of total N-cycle gene copies versus hydrogen index (HI); right panel: square root of total N-cycle gene copies versus organic carbon content (C_org_). Points are colored and shaped by site. Spearman correlation coefficients (ρ) and p-values are shown for each relationship.

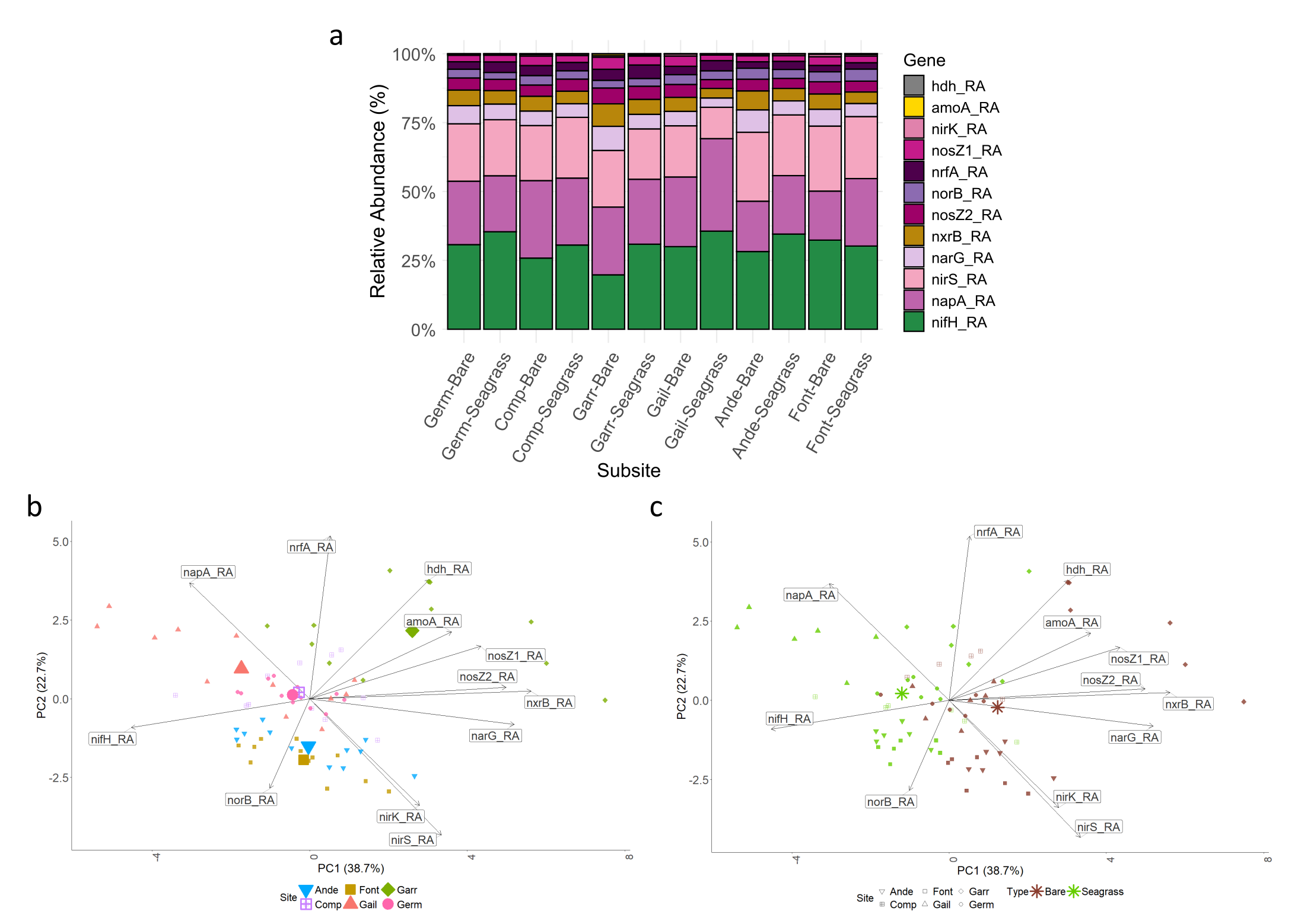

**Figure S3.** Targeted metagenomic profiling of N-cycling genes across sediment types and sites. (a) Relative abundance of the 12 targeted N-cycling genes across subsites, shown as stacked bar plots. Gene abundances are expressed as relative proportions within the total pool of targeted N-cycling genes for each sample. (b) PCA based on the relative abundance of the 12 N-cycling genes. Colors and point shapes indicate sampling sites. Vectors indicate the contribution of each gene to the ordination axes. (c) PCA based on the same dataset, with points colored according to sediment type (Seagrass vs. Bare sediment) and shapes representing site.

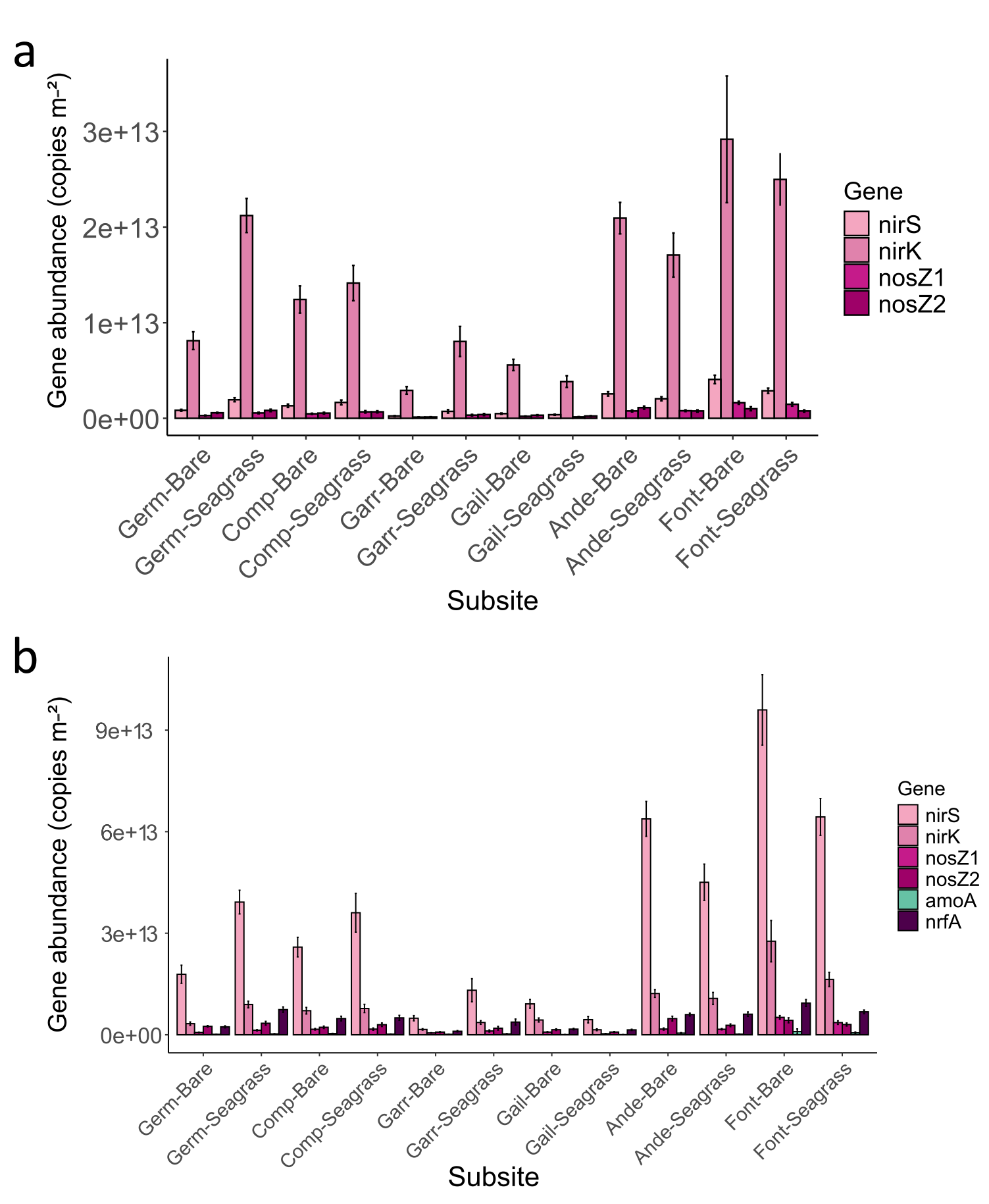

**Figure S4.** Comparison of gene abundances quantified by qPCR and estimated from targeted metagenomic data. (a) shows gene abundances quantified by qPCR. (b) shows absolute abundances calculated by multiplying qPCR copy numbers by the relative abundance obtained from targeted metagenomic data (both in copies m⁻²).

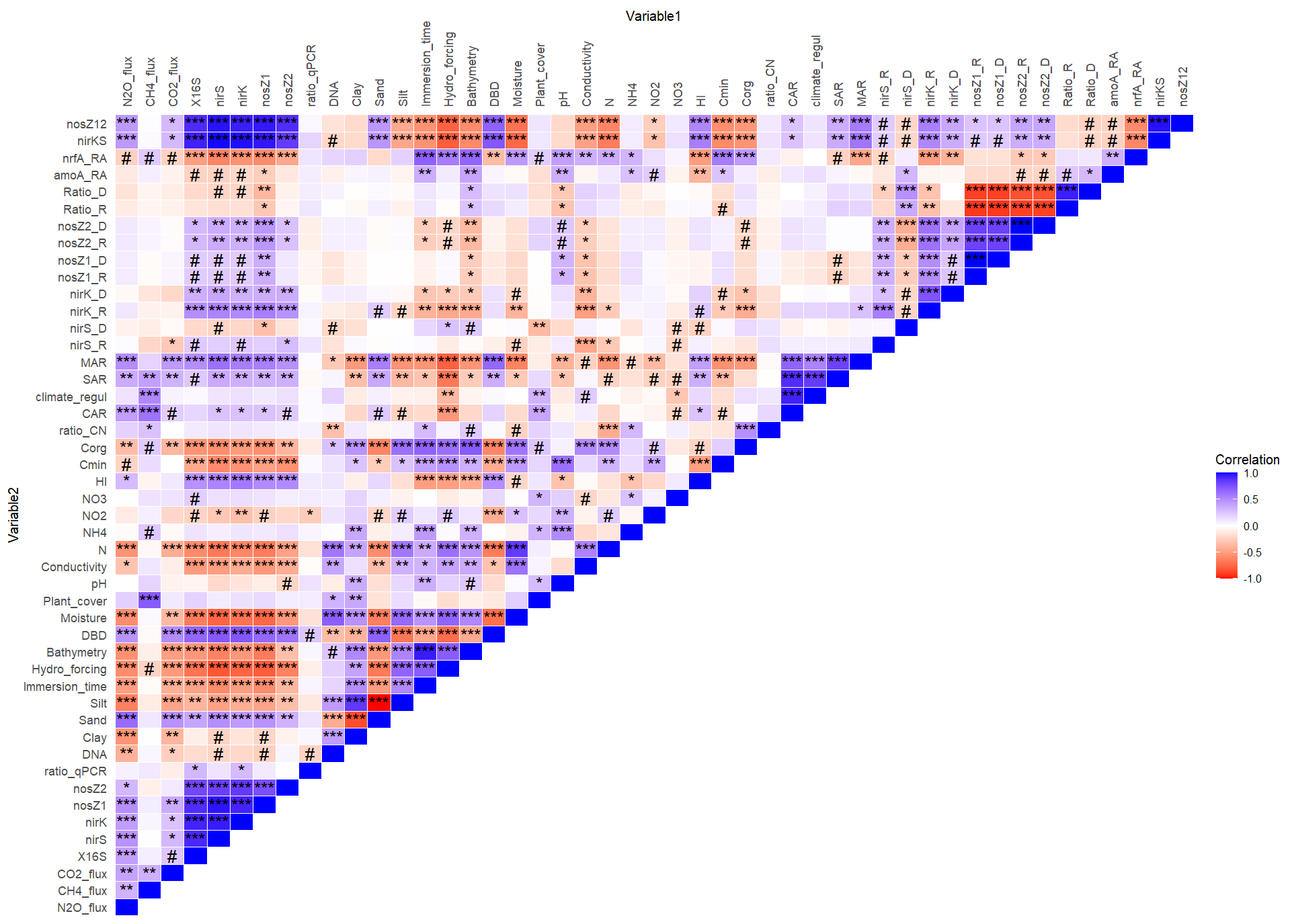

**Figure S5.** Correlation matrix of physical, chemical, microbial, and GHG variables across all sampling sites in Arcachon Bay**.** Significant correlations are indicated with # (p < 0.1), * (p < 0.05), ** (p < 0.01), *** (p < 0.001), and non-significant correlations are left blank.

a *nirS*

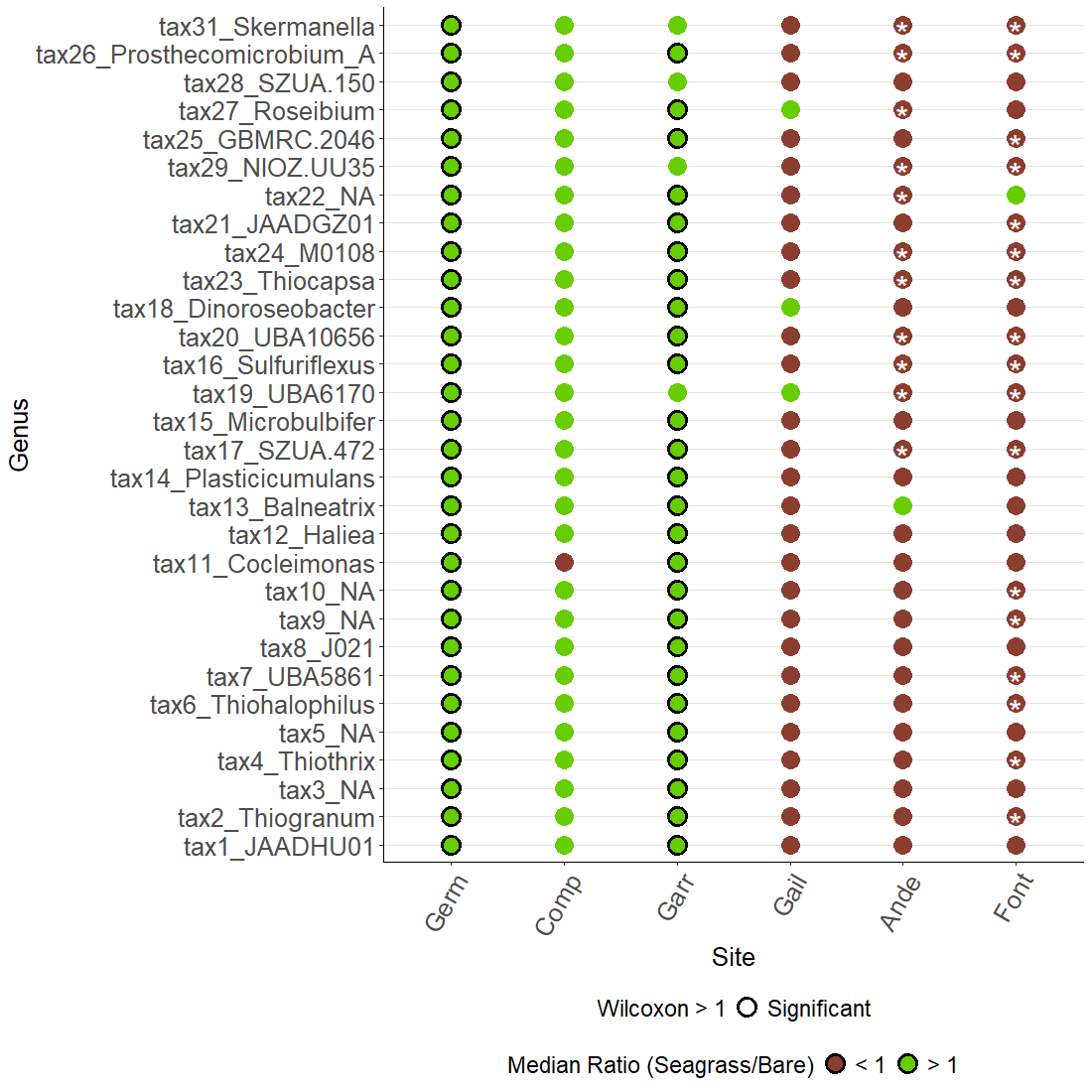

b *nirK*

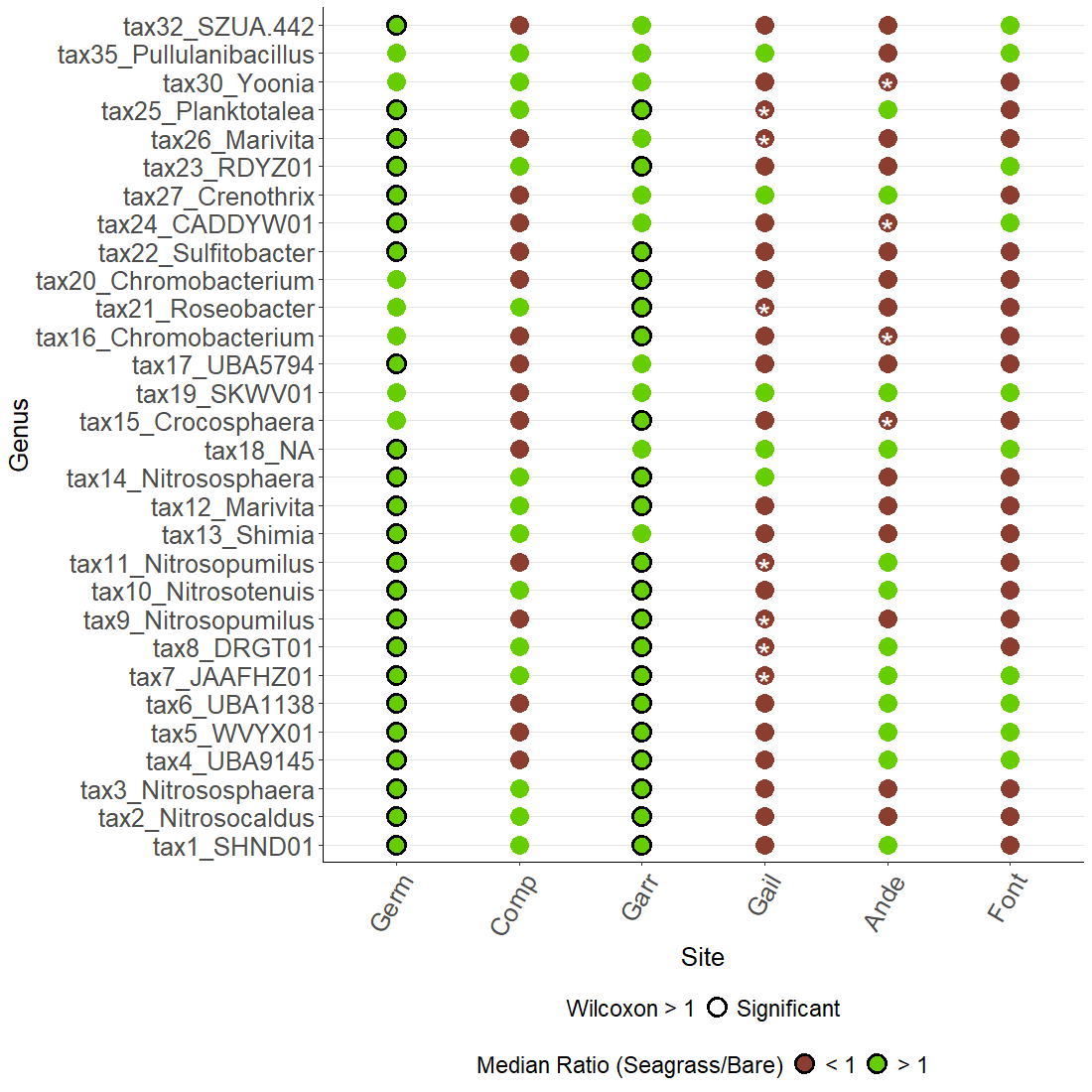

c *nosZ1*

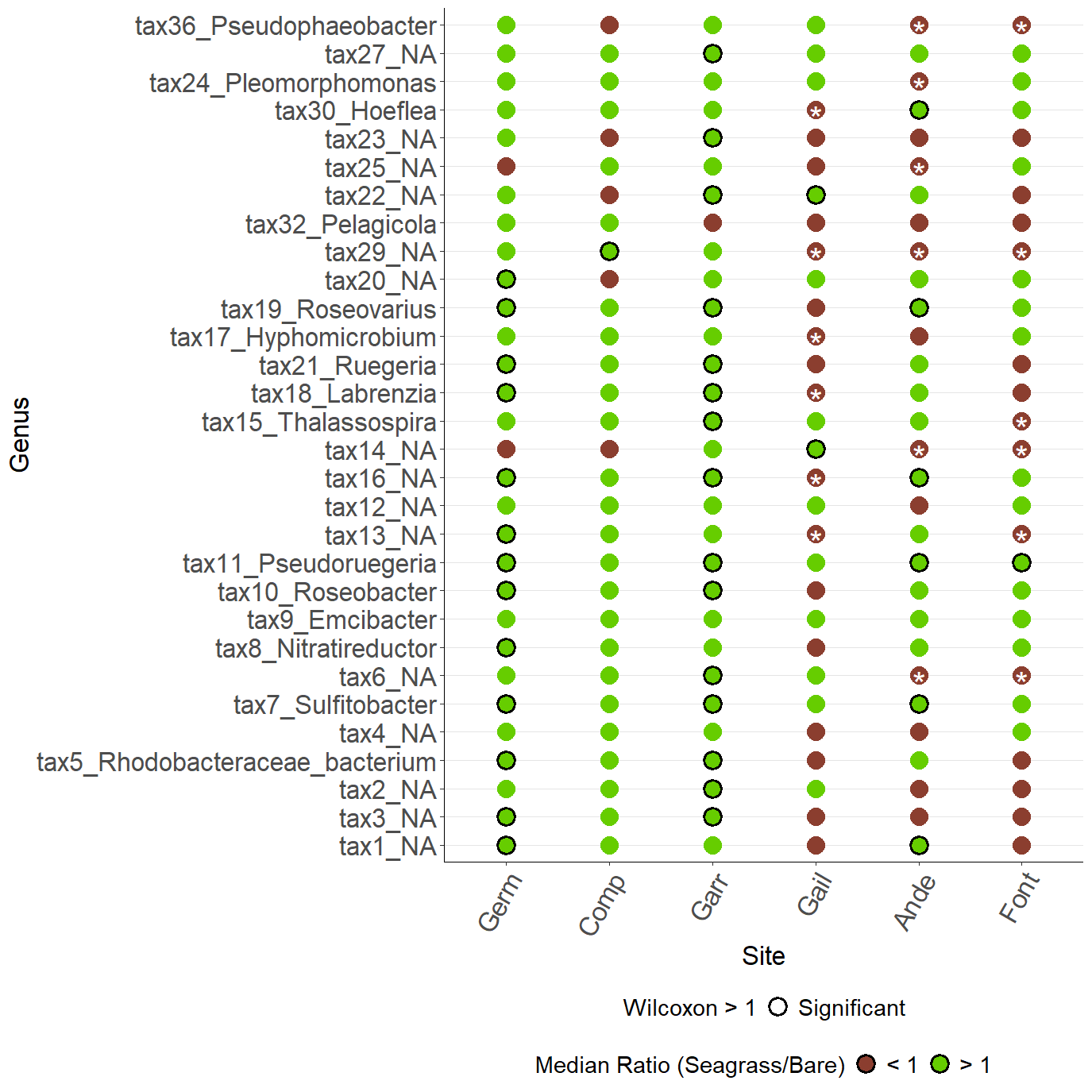

d *nosZ2*

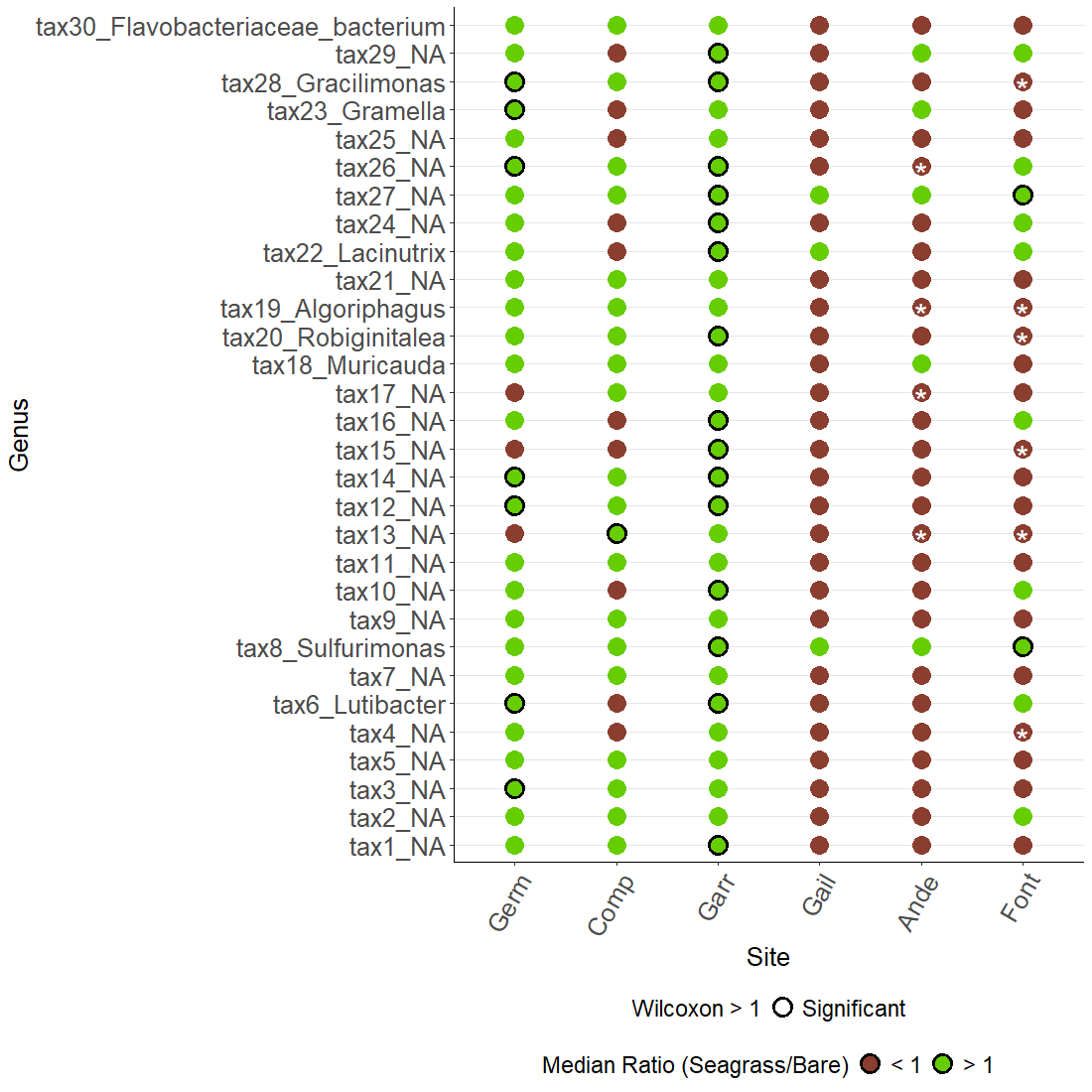

**Figure S6.** Differential abundance and relationships with N₂O flux for N₂O-producing and N₂O-consuming genera across sediment type in Arcachon Bay. (a–d) Genera associated with the *nirS* (a), *nirK* (b), *nosZ1* (c), and *nosZ2* (d) genes. For each genus, the panel displays median abundance ratios: Seagrass/Bare, color-coded (green: ratio > 1, Seagrass affinity; brown: ratio < 1, Bare affinity). Significant differences (Wilcoxon test, P < 0.05) are indicated by a black circle for seagrass preference and a white star within brown point for Bare preference. Sites are ordered by increasing N₂O flux in seagrass sediments.

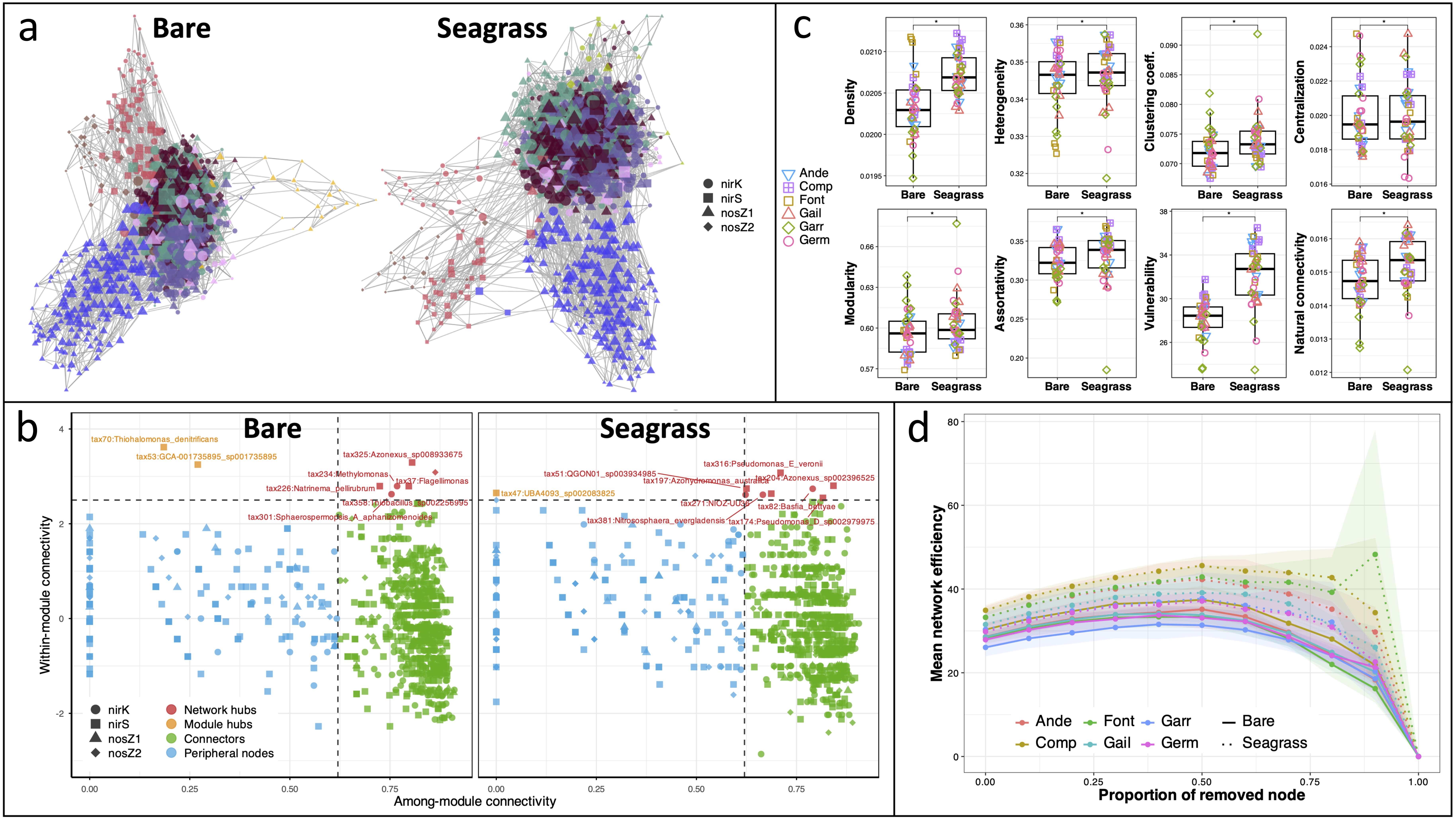

**Figure S7.** Comparison of microbial co-occurrence network topology between Bare and Seagrass sediment communities. (**a)** Co-occurrence networks constructed for the Bare and Seagrass sites. Nodes represent microbial taxa, with shapes indicating the associated N₂O-cycling functional genes (*nirK*, *nirS*, *nosZ1*, and *nosZ2*). Node colors indicate the module membership of each taxon. Edges represent significant co-occurrence relationships between taxa. **(b)** Topological roles of nodes based on within-module connectivity (*Zi*) and among-module connectivity (*Pi*). Dashed lines indicate the thresholds (*Zi* = 2.5 and *Pi* = 0.62) used to classify nodes as peripheral nodes, connectors, module hubs, or network hubs. Taxa identified as connectors or hubs are annotated. (**c)** Comparison of network topological properties between Bare and Seagrass networks, including density, heterogeneity, clustering coefficient, centralization, modularity, assortativity, vulnerability, and natural connectivity. Points represent individual sampling sites, boxplots summarize the distribution of network metrics, and asterisks indicate significant differences between Bare and Seagrass networks (ns, not significant; ****, P < 0.0001). (**d)** Network robustness assessed by progressive low-degree node removal. Mean network efficiency was calculated from 20 simulations and averaged between 25% and 90% node removal. Colored lines represent the six sampling sites, whereas solid and dotted lines correspond to the Bare and Seagrass networks, respectively. Shaded areas indicate the variability among simulations.
